# MT5-MMP controls APP and β-CTF/C99 metabolism through proteolytic-dependent and -independent mechanisms relevant for Alzheimer’s disease

**DOI:** 10.1101/2020.09.01.258665

**Authors:** Laura García-González, Jean-Michel Paumier, Laurence Louis, Dominika Pilat, Anne Bernard, Delphine Stephan, Nicolas Jullien, Frédéric Checler, Emmanuel Nivet, Michel Khrestchatisky, Kévin Baranger, Santiago Rivera

## Abstract

We previously discovered the implication of membrane-type 5-matrix metalloproteinase (MT5-MMP) in Alzheimer’s disease AD pathogenesis. Here we shed new light on pathogenic mechanisms by which MT5-MMP controls APP processing and the fate of amyloid beta peptide (Aβ), its precursor C99 and C83. We found in HEK carrying the *APP* Swedish familial mutation (HEKswe) that MT5-MMP-mediated processing of APP that releases the soluble 95 kDa form (sAPP95), was hampered by the removal of the C-terminal non-catalytic domains of MT5-MMP. Catalytically inactive MT5-MMP variants increased the levels of Aβ and promoted APP/C99 sorting in the endo-lysosomal system. We found interaction of C99 with the C-terminal portion of MT5-MMP, the deletion of which caused a strong degradation of C99 by the proteasome, preventing Aβ accumulation. These findings reveal novel mechanisms for MT5-MMP control of APP metabolism and C99 fate involving proteolytic and non-proteolytic actions mainly mediated by the C-terminal part of the proteinase.

## INTRODUCTION

Deciphering the pathophysiological mechanisms of Alzheimer’s disease (AD) on which to base the design of efficient therapies represents a major scientific, societal and economic challenge. Amyloid beta peptide (Aβ) resulting from the cleavage of the amyloid beta precursor protein (APP), has long been considered to be the major pathogenic factor in AD, but other neurotoxic APP fragments are emerging in a context of more complex proteolytic processing of APP than expected (reviewed in [1-3]). APP cleavage by β-secretase generates a transmembrane β-C-terminal fragment of 99 amino acids (β-CTF/C99), which is then cleaved by γ-secretase to release monomeric forms of Aβ. Cytotoxic assembly and accumulation of monomers into oligomers and fibrils is the cornerstone of the amyloid cascade hypothesis of AD [4]. Although much less studied, the toxicity of the immediate precursor of Aβ, C99, has also been documented [5-10]. Thus, C99 accumulation occurs in the brains of the 3xTg and 5xFAD mouse models of AD, where it precedes that of Aβ [8,11,12,10]. C99 also accumulates in fibroblasts and brains of AD patients [13,14], as well as in human neurons derived from induced pluripotent stem cell (iPS) with AD familial mutations, resulting in pathogenic endosomal dysfunction and altered subcellular trafficking [15]. Taken together, these data are consistent with the contribution of C99 to the pathophysiology of AD and justify the study of APP metabolism beyond Aβ, including the role of proteinases involved in its regulation.

Membrane-type 5-matrix metalloproteinase (MT5-MMP, also known as MMP-24 or η-secretase) is one of the newly discovered APP-cleaving enzymes [2]. Primarily expressed in neural cells [16,17], MT5-MMP contributes to physiological and pathological processes in the nervous system [18,19]. Its colocalization with amyloid plaques in the brain of AD patients first suggested a possible implication in the disease [17]. We confirmed this in 5xFAD mice, where MT5-MMP deletion resulted in a strong reduction of C99 and Aβ content, reduced levels of neuroinflammatory mediators, and preservation of long-term potentiation (LTP) and cognition in the early stages of the disease. These effects were observed without changes in β- or γ-secretase activities [12,20]. In addition, MT5-MMP could release a soluble N-terminal fragment of ∼95 kDa (sAPP95) and increase Aβ and C99 levels in HEKswe [12,20]. Importantly, sAPP95 levels decreased in the brains of MT5-MMP-deficient 5xFAD mice, identifying APP as a genuine *in vivo* substrate for the proteinase [12,20]. The corresponding η-cleavage site was reported, along with the ability of MT5-MMP to generate a synaptotoxic C-terminal APP fragment (*i*.*e*., Aη-α) in concert with ADAM10 [21]. In conclusion, MT5-MMP promotes the accumulation of various toxic APP metabolites and associated functional alterations, while these are prevented by proteinase suppression.

In this study, we sought to better understand the mechanisms that govern the functional interaction(s) between MT5-MMP and APP and that significantly influence the metabolism of the latter. We used molecular and cell biology, as well as structure-function approaches to reveal novel proteolytic and, more unusually, non-proteolytic actions of specific domains of MT5-MMP that determine the levels of sAPP95, C99 and Aβ. Our work provides key new insights into the molecular mechanisms that support the pathogenic actions of MT5-MMP in AD and paves the way for therapeutic strategies to target this proteinase.

## METHODS

### Reagents

MMP-2 inhibitor III and BACE inhibitor IV (C3) (both from Millipore, Molsheim, France) and the γ-secretase inhibitor DAPT (Tocris, Bio-Techne, Lille, France) were used at 10 μM. The proteasome inhibitor MG132 was used at 5 μM (Enzo Life Science, Lyon, France) and bafilomycin A1 (Sigma-Aldrich, Saint-Quentin Fallavier, France) was used at 50 nM. RxP03, which targets MMPs and spares adamalysins, was used at 10 μM [22-24]. All the restriction enzymes were purchased from New England Biolabs (Evry, France). All the chemical reagents used in this study were from analytic grade and purchased from Sigma-Aldrich, and all the products/media required for cell culture were from Thermo Fischer Scientific (Villebon-sur-Yvette, France), unless otherwise stated.

### Plasmid constructs

cDNAs encoding *MMP24* (MT5-MMP), *MMP14* (MT1-MMP), *TIMP1* (TIMP-1) and *TIMP2* (TIMP-2) were amplified by PCR from P15 C57Bl6 mice cerebellum mRNA. MT5-MMP (MT5/GFP), active and inactive MT1-MMP (MT1/GFP and MT1Δ/GFP), TIMP-1/RFP and TIMP-2/RFP were cloned and expressed as previously reported [20,24]. To generate inactive MT5-MMP (MT5Δ/GFP), the Quick Change Lighting MultiSite-directed Mutagenesis kit (Agilent Technologies, Les Ulis, France) was used and the primers were as follows: *MT5*Δ (*E256Q*)*For* CTG CTG GCC GTG CAT CAA CTG GGC CAT GCA CTG and *MT5*Δ *(E256Q)Rev* CAG TGC ATG GCC CAG TTG ATG CAC GGC CAC CAG. All the genes were cloned in pEGFP-N1 or pDSRed II-N1 [20,24]. Plasmids within a pcDNA3.1 backbone encoding full length active (MT5FL) and inactive (MT5Δ) MT5-MMP and the truncated MT5-MMP constructs (CAT^-^, HPX^-^, EXT^-^, TM/IC^-^ and IC^-^) were synthesized and produced by GeneArt (Thermo Fisher Scientific). All these constructs were fused with a FlagM2 tag on their N-terminal part (Fig. 4A). Full length MT1-MMP (MT1FL) and the truncated constructs (CAT^-^, HPX^-^, EXT^-^ and IC^-^) were cloned in pcDNA3.1 vector and kindly provided by Pr. Motoharu Seiki and Pr. Takeharu Sakamoto [25]. The plasmid that encodes human C99 (pcDNA3.1-C99) was previously described [26]. We used the pcDNA3.1-C99 construct as a template to generate a pcDNA3.1-C99 with HA tag on the N-terminal (HA-C99), using the SLIC method [27]. All the plasmids were amplified in *E. coli* DH5α (Thermo Fisher Scientific) and purified using the NucleoBond Xtra Midi Plus EF (Macherey-Nagel, Hoerdt, France), according to the manufacturer’s recommendations.

**Figure 1:**
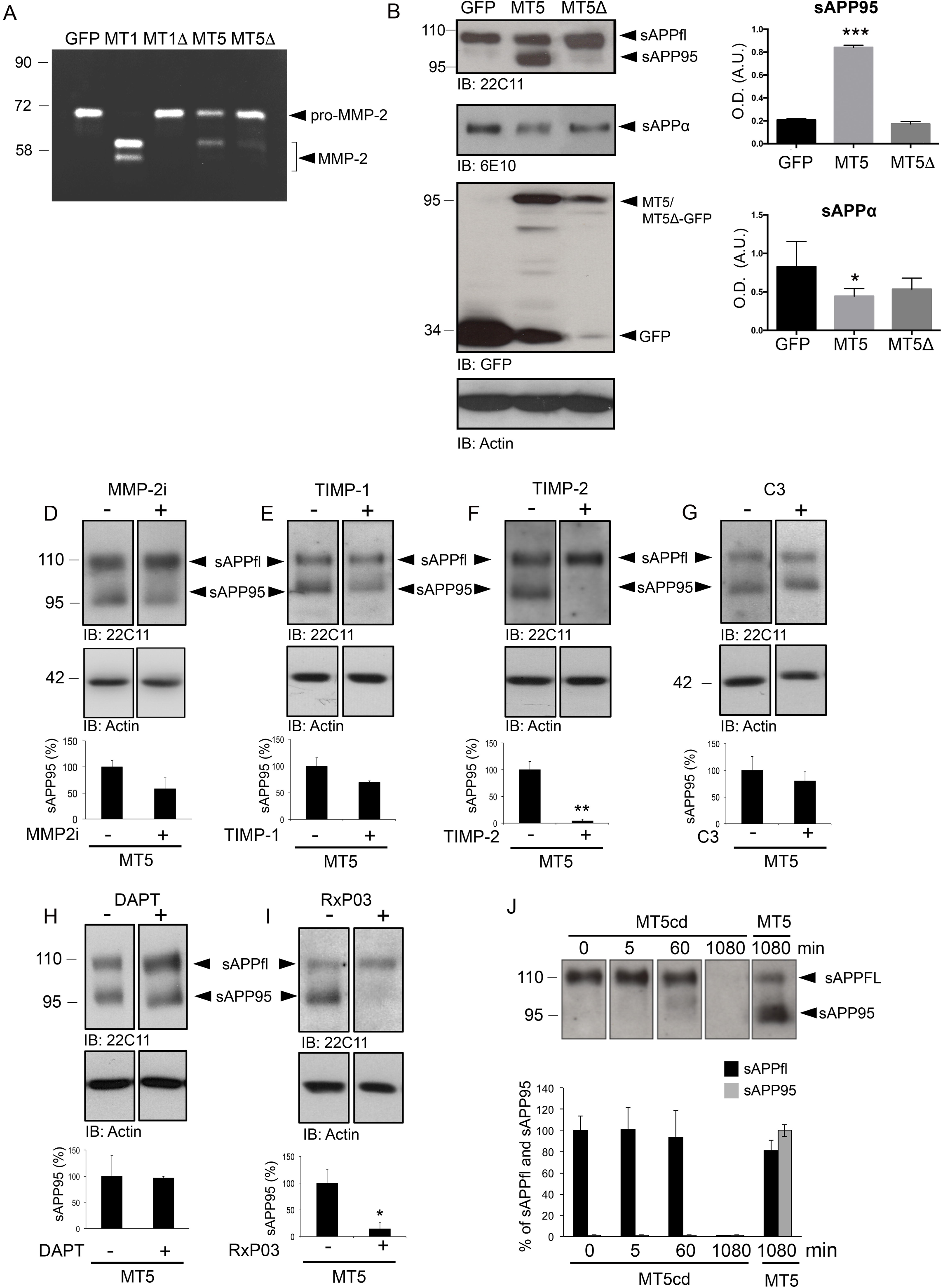
MT5-MMP-mediated generation of sAPP95 is dependent of its catalytic activity. **A**. Gelatin zymography on supernatants of HEKswe cells 48 h after transfection with GFP control or active (MT1) or inactive (MT1Δ) MT1-MMP/GFP, and active (MT5) or inactive (MT5Δ) MT5-MMP/GFP constructs. Note that MT5 is less efficient for MMP-2 activation than MT1. **B**. Representative western blots (WB) of sAPPFL and sAPP95 immunoreactivity in HEKswe supernatants using anti-APP 22C11 (top), and WB of sAPPα (2^nd^ from top) using anti-Aβ 6E10 antibodies. The 3^rd^ from top panel represents WB of GFP and active and inactive MT5-MMP/GFP expression in cell lysates 48 h after transfection using anti-GFP antibody. The bottom panel represents actin-loading controls. **C**. Histograms representing the actin-normalized quantifications for sAPP95 (top) and sAPPα (bottom)**D, E, F, G, H** and **I**. WB and the corresponding actin-normalized quantifications of sAPPFL and sAPP95 levels using the anti-APP 22C11 antibody in the supernatants of HEKswe cells after transfection with active MT5-MMP/GFP (MT5) in the presence or not of 10 μM of MMP-2 selective inhibitor III (MMP-2i) (D), TIMP-1/RFP (co-transfection) (E), TIMP-2/RFP (co-transfection) (F), 10 μM of C3 (G), 10 μM of DAPT (H) and 10 μM of RXP03 (I). **J**. WB and the corresponding quantifications of immunoreactive bands for anti-APP 22C11 antibody representing sAPPFL and sAPP95 in cell-free conditioned media from control GFP-transfected HEKswe incubated at 37°C up to 18 h (1080 min) with the recombinant MT5-MMP catalytic domain (MT5cd). The right lane represents a control of sAPPFL and sAPP95 production obtained from the supernatants of HEKswe cells transfected with MT5 that were further incubated with vehicle control for 1080 min h. Note that MT5cd was unable to generate sAPP95 *in vitro*. Values are the mean +/-SEM of at least four independent cultures. * p<0.05 and *** p<0.001 *vs* GFP untreated group. ANOVA followed by *post hoc* Fisher’s LSD test (**C**) and Student t-test (**D-I**).

**Figure 2.**
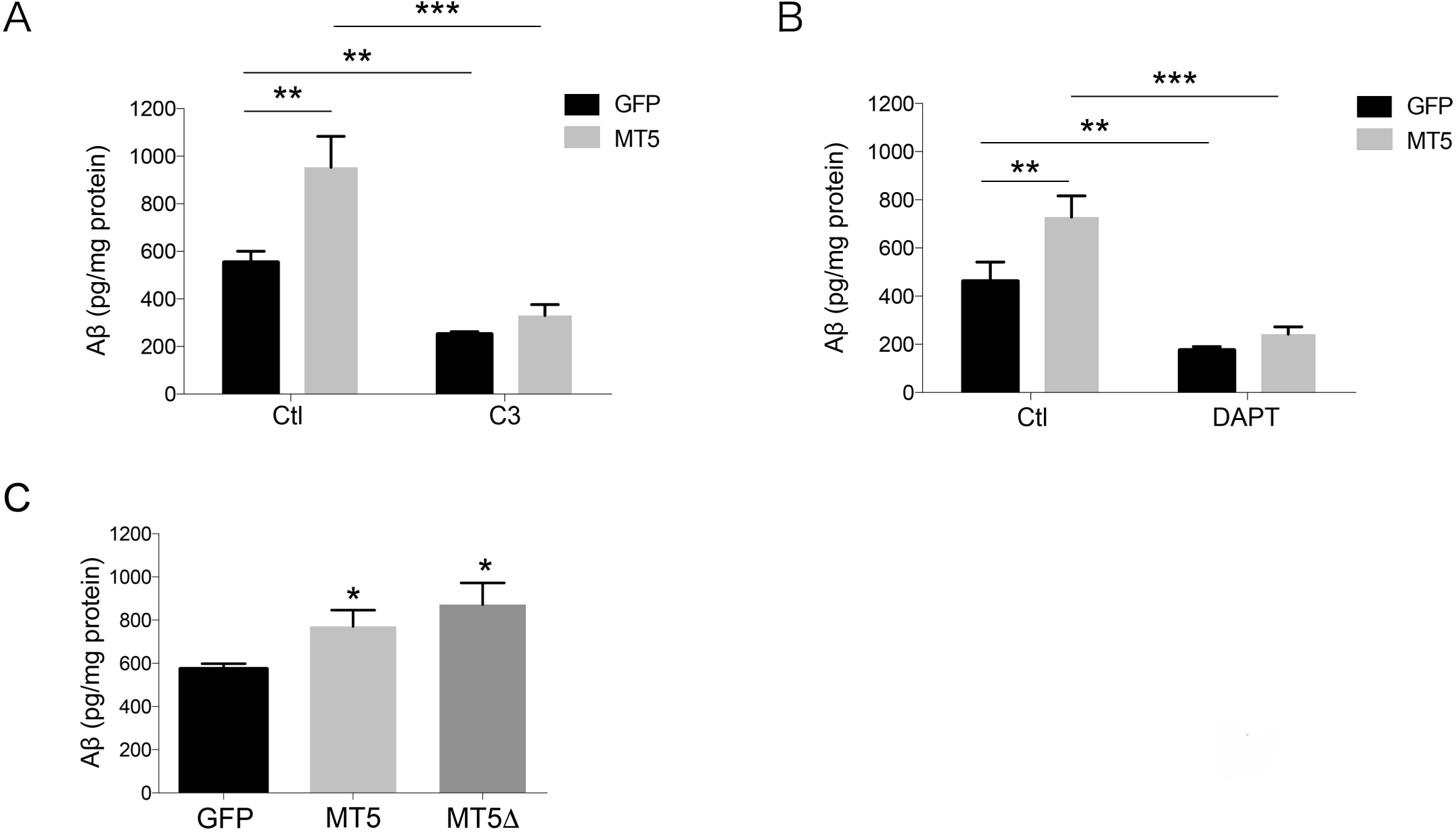
β- and γ-secretase mediate the upregulation of Aβ levels by MT5-MMP and inactive MT5-MMP is also pro-amyloidogenic. **A, B**. Histograms showing ELISA quantification of Aβ40 levels (pg/mg protein) in the supernatants of HEKswe cultures (n= 4-6) 48 h after transfection with GFP control and active MT5-MMP/GFP (MT5) in the presence or not of 10 μM of C3 (A) and 10 μM of DAPT (B). **C**. Histogram showing ELISA quantification of Aβ40 levels (pg/mg protein) in the supernatants of HEKswe cultures (n= 4-9) 48 h after transfection with GFP control, active (MT5) or inactive (MT5Δ) MT5-MMP/GFP constructs. Note that MT5Δ is as effective as MT5 in stimulating Aβ accumulation. Values are the mean of the indicated independent cultures. * p<0.05, ** p<0.01 and *** p<0.001 *vs* GFP untreated group. ANOVA followed by *post hoc* Fisher’s LSD test.

**Figure 3.**
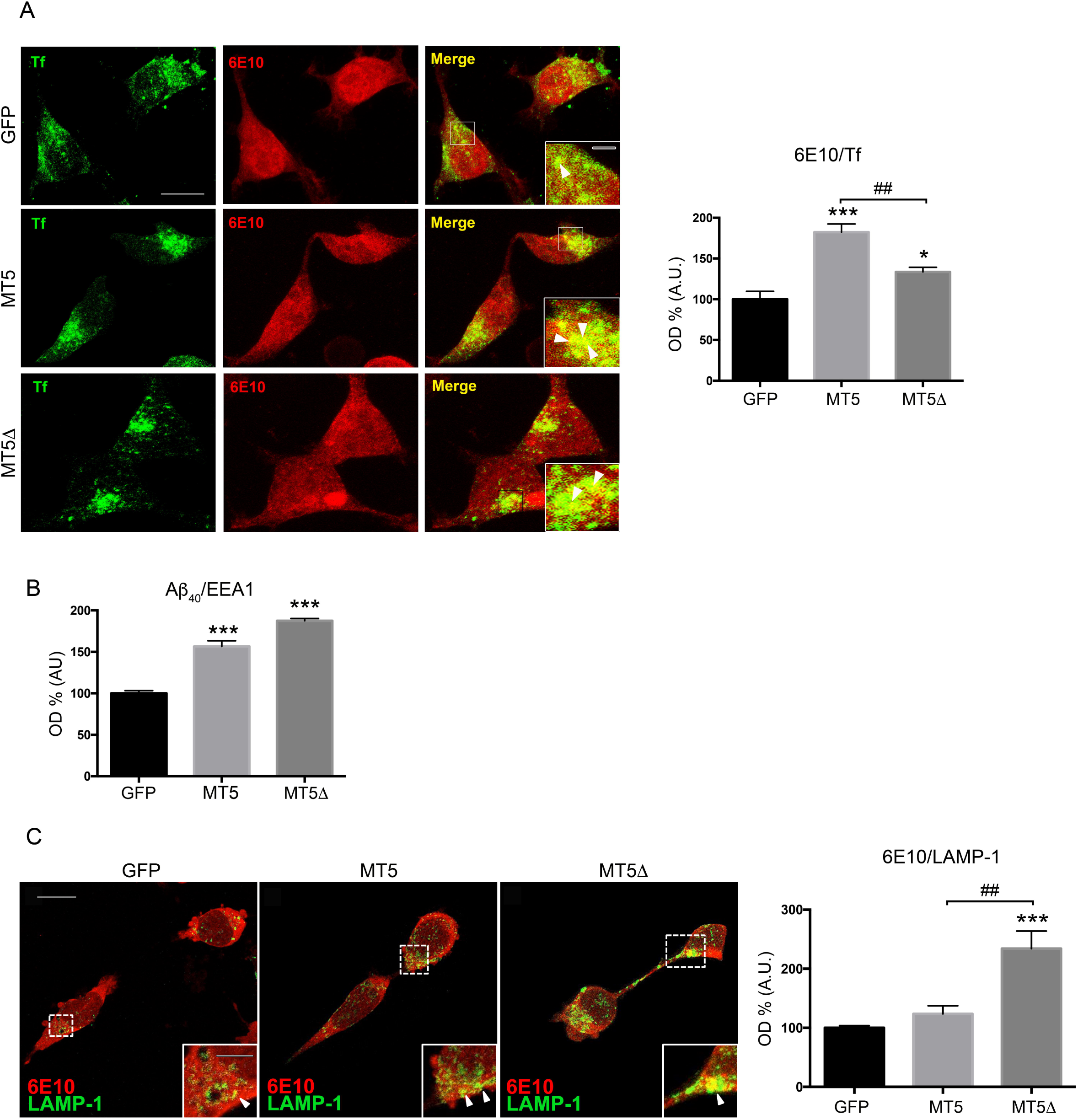
MT5-MMP modulation promotes the distribution of APP/Aβ in early endosomes and lysosomes. **A**. Confocal micrographs showing anti-6E10^+^ immunoreactivity (red) in HEKswe cells incubated for 30 min at 37°C with transferrin-AlexaFluor^647^ (green; Tf) to label early endosomes, 48 h after transfection with constructs coding for GFP control (top panels), active MT5-MMP/GFP (MT5) (middle panels) or inactive MT5Δ-MMP/GFP (MT5Δ) (bottom panels). Arrowheads in the inset of the merge panel illustrate areas of colocalization between 6E10^+^ and Tf^+^ signals (yellow). The histogram shows the % of colocalization in arbitrary units (A.U.) between 6E10^+^ and Tf^+^ with respect to GFP control. **B**. Histograms showing the % of colocalization between Aβ40^+^ and EEA1^+^ (early endosomes) signals in HEKswe cells transfected with GFP, MT5 or MT5Δ constructs. **C**. Confocal micrographs showing anti-6E10^+^ (red) and anti-LAMP-1^+^ immunoreactivity (green) in HEKswe cells 48 h after transfection with either GFP, MT5 or inactive MT5Δ constructs. Arrowheads in the inset of the merge panel illustrate areas of colocalization between 6E10^+^ and LAMP-1^+^ signals (yellow). The histogram shows the % of colocalization between 6E10^+^ and LAMP-1^+^ signals with respect to GFP control. Values represent the mean +/-SEM of four independent cultures. ** p<0.01 and *** p<0.01 *vs* GFP. ^##^ p<0.01 *vs* MT5. ANOVA followed by *post hoc* Fisher’s LSD test. Scale bars 10 µm. Scale bar in the insets, 3 µm.

**Figure 4.**
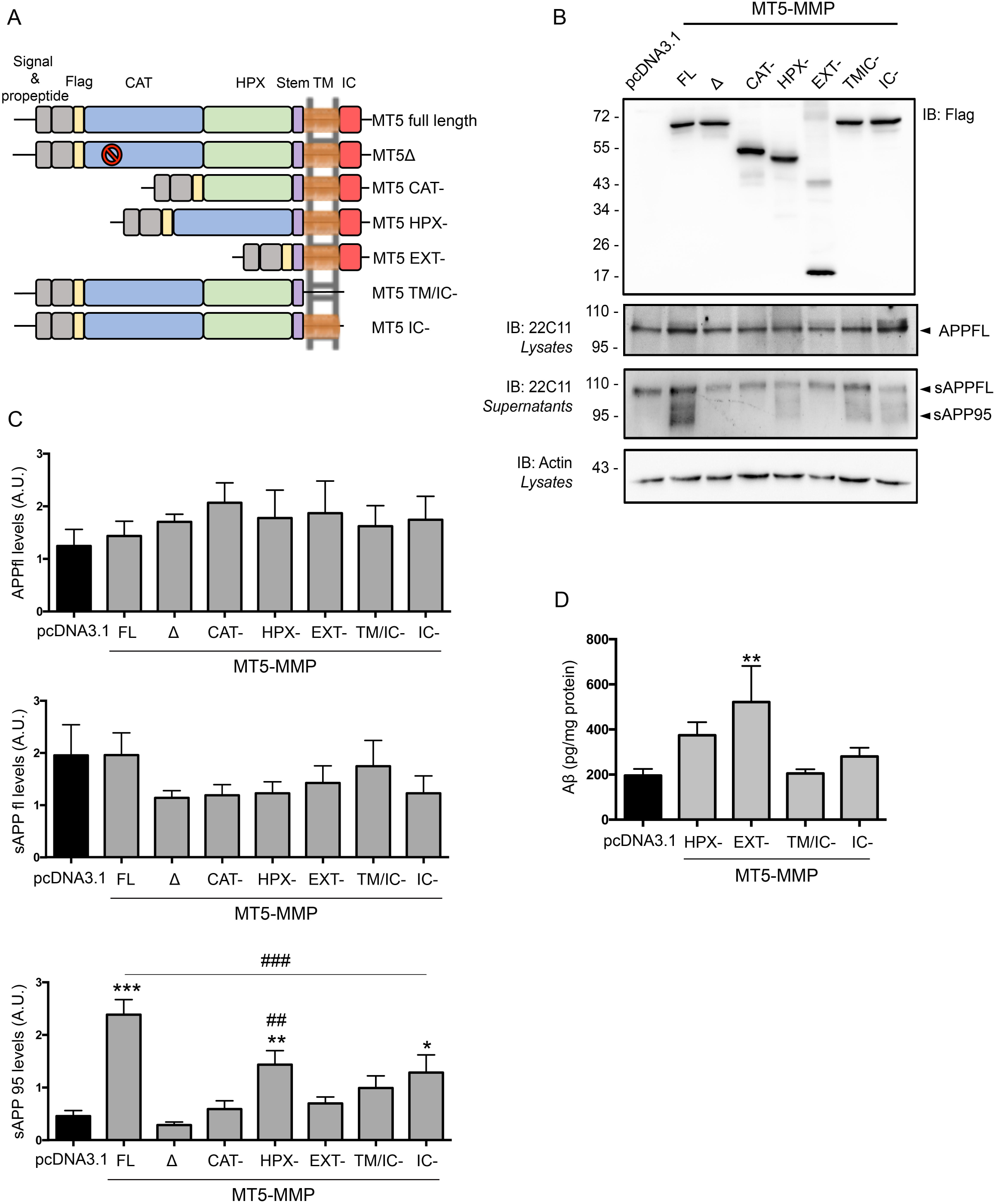
MT5-MMP domains differently affect sAPP95 and A β production. **A**. Schematic representation of different MT5-MMP variants tagged with flagM2 on the N-ter, including full length active (MT5 FL) and inactive (MT5Δ) MT5-MMP, as well as truncated MT5-MMP variants without catalytic (CAT-), hemopexin (HPX-), extracellular (EXT-), transmembrane/intracytoplasmic (TM/IC-) and intracytoplasmic (IC-) domains. **B**. WB representing these MT5-MMP variants and pcDNA3.1 control plasmid as control 48 h after transient transfection of HEKswe cells, using an anti-FlagM2 antibody (1^st^ panel top). Total full length APP (APPFL) from cell lysates (2^nd^ panel) and sAPPFL and sAPP95 in cell supernatants (3^rd^ panel), using the anti-APP 22C11 antibody. Actin-loading controls are represented in the 4^th^ panel. **C, D** and **E**. Histograms showing actin-normalized quantifications of APPFL in cell lysates (C), sAPPFL (D) and sAPP95 (E) in cell supernatants. **F**. Histogram showing ELISA quantification of Aβ40 levels (pg/mg protein) in the supernatants of HEKswe cultures (n=4) 48 h after transfection with pcDNA3.1, HPX-, EXT-, TM/IC- and IC-constructs. Note that only EXT- increases Aβ levels with respect to pcDNA3.1, TM/IC- and IC-groups. Values represent the mean +/-SEM of 5 independent cultures. * p<0.05, ** p<0.01, *** p<0.005 *vs* pcDNA3.1. ^##^ p<0.01 and ^###^ p<0.001 *vs* MT5 FL. ANOVA followed by *post hoc* Fisher’s LSD test.

### Transfection of HEK cells

GFP, MT5/GFP and MT5Δ/GFP encoding plasmids were transfected in HEK cells stably expressing human *APP* with the Swedish mutation (HEKswe) under the control of the CMV promoter, as previously described [12,20,24]. Cells were plated to 10^6^ cells/well for 24 h in 6-well plates in DMEM Glutamax, 10% FBS, 1% penicillin/streptomycin (P/S). Cells were transfected with a final concentration of 1 μg of plasmids, using Jet Pei transfection reagent (Ozyme, Saint-Quentin en Yvelines, France). Sixteen hours after transfection, the medium was replaced with OptiMEM (serum-free medium) containing 1% Insulin-Transferrin-Selenium (ITS) and cells were then allowed to secrete for 48 h.

Plasmids (0.5 μg/plasmid) encoding MT5FL, MT5Δ and the different MT5-MMP truncated variants CAT^-^, HPX^-^, EXT^-^, TM/IC^-^, IC^-^ or those derived from MT1-MMP *i*.*e*., MT1FL, CAT^-^, HPX^-^, EXT^-^, IC^-^ were co-transfected with C99 or HA-C99 (0.5 μg/plasmid) in HEK cells using the Jet Pei transfection reagent. Cells were plated to 10^6^ cells/well for 24 h in 6-well plates in DMEM Glutamax, 10% FBS, 1% P/S. Sixteen hours after transfection, the medium was replaced with OptiMEM (serum-free medium) containing 1% ITS and cells were then treated for 48 h in the presence or not of DAPT, MG132 and bafilomycin A1. DMSO (0.01%) was used as control vehicle for these drugs.

### Reverse transcription-quantitative polymerase chain reaction (RT-qPCR)

Total RNA was extracted from HEKswe cells using the Nucleospin RNA kit (Macherey-Nagel), according to the manufacturer’s recommendations. Single-stranded cDNA was synthesized from 1 μg of RNA using High-Capacity RNA to cDNA™ kit (Thermo Fisher Scientific) suitable for quantitative PCR. Twenty-five ng of cDNA were submitted to qPCR reaction using the Fast Real-Time PCR System (Thermo Fisher Scientific). For each experiment, cDNA samples were analysed in duplicate and relative gene expression was obtained using the comparative 2^-(ΔΔCt)^ method after normalization to the *Gapdh* housekeeping gene [11,24]. All the probes were purchased from Thermo Fisher Scientific (Table 1).

**Table 1.** Human TaqMan probes used for quantitative PCR analysis.

| **Gene name** | **Gene description** | **Probes ID** |
| --- | --- | --- |
| *MMP2* | Matrix metalloproteinase 2 | Hs01548727_m1 |
| *MMP9* | Matrix metalloproteinase 9 | Hs00234579_m1 |
| *MMP24* | Membrane-type 5 matrix metalloproteinase | Hs01084645_m1 |
| *MMP14* | Membrane-type 1 matrix metalloproteinase | Hs01037009_g1 |
| *TIMP1* | Tissue inhibitor of matrix metalloproteinase 1 | Hs00171558_m1 |
| *BACE1* | Beta-site APP-cleaving enzyme-1 | Hs01121195_m1 |
| *PSEN1* | Presenilin 1 | Hs00997789_m1 |
| *PSEN2* | Presenilin 2 | Hs01577197_m1 |
| *PSENEN* | Presenilin enhancer 2 homolog | Hs00708570_s1 |
| *APH1A* | Aph1a gamma secretase subunit | Hs01046142_g1 |
| *NCSTN* | Nicastrin | Hs00950933_m1 |
| *MME* | Membrane metallo-endopeptidase<br>(*neprilysin*) | Hs00153510_m1 |
| *CTSD* | Cathepsin D | Hs00157205_m1 |
| *APP* | Amyloid beta (A4) precursor protein | Hs00169098_m1 |
| *GAPDH* | Glyceraldehyde-3-phosphate dehydrogenase | Hs02758991_g1 |

### Cell viability

Cell viability was evaluated using the 3-(4,5-Dimethylthiazol-2yl)-2,5-diphenyl tetrazolium bromide (MTT) assay (Sigma-Aldrich), which measures mitochondrial activity in living cells. Data were calculated as the percentage of living cells = (transfected cell OD_550_/control cell OD_550_) x 100.

### Western blot, *in vitro* APP cleavage and immunoprecipitation assay

Supernatants from HEKswe or HEK cells were collected 48 h after transfection with the different MT5-, MT1-MMP and C99 constructs and centrifuged at 300 x *g* for 5 min at 4°C. Cells were scraped in RIPA buffer (Sigma-Aldrich) containing proteinase inhibitors (Millipore) and sonicated. Protein concentration was determined using a Bio-Rad *DC*™ protein assay kit (Bio-Rad, Marnes-La-Coquette, France). Proteins (30 µg) were loaded on 10% to 15% SDS-PAGE gels, low molecular weight gels Tris-Tricine gels or pre-casted gels 4-20% (Thermo Fisher Scientific), and transferred onto nitrocellulose membranes (GE Healthcare, Dutscher, Brumath France). After blocking, membranes were probed with the following primary antibodies: APP 22C11 (1/1000, Millipore), APP-CTF (1/1000, Sigma-Aldrich), APP 6E10 (1/1000, Ozyme), anti-HA (1/1000, Roche Diagnostics, Meylan, France), anti-FlagM2 (1/1000, Sigma-Aldrich), anti-MT1-MMP (Abcam, Cambridge, UK), anti-GFP (1/2000, Roche Diagnostics), anti-actin (1/5000, Sigma-Aldrich). Then, the appropriate HRP-conjugated secondary IgG antibodies were used (Jackson Immunoresearch). To test whether sAPP95 may result from proteolysis of full length sAPP (sAPPFL) by MT5-MMP, 2 nM of recombinant catalytic domain of MT5-MMP (MT5cd, Enzo Life Science) was incubated with supernatants from GFP-transfected HEKswe at 37°C in activity buffer (50 mM Tris-HCl pH 7.5, 150 mM NaCl, 5 mM CaCl_2_, 0.025% Brij35) for 5 min, 1 h and 18 h. Samples were loaded on 10% SDS-PAGE gels and incubated with 22C11 antibody as described above.

For the immunoprecipitation (IP) assay, 250 μg of protein (cell lysates) in 500 μL of RIPA buffer were incubated overnight at 4°C with unspecific mouse/rabbit/rat IgG antibodies (Jackson Immunoresearch) or rat anti-HA-tag (Roche Diagnostics) or mouse anti-FlagM2 and rabbit anti-APP-CTF (Sigma-Aldrich) at 4 ng/μL and then pulled-down for 2 h using protein G-coupled dynabeads (50 μL, Thermo Fisher Scientific). After washing 3 times with 0.3 M NaCl solution, samples were denatured and subjected to western blot (WB) using APP-CTF, anti-FlagM2 or anti-HA antibodies, and the corresponding horseradish peroxidase-conjugated secondary IgG antibodies (Jackson Immunoresearch). The EasyBlocker kit was used to limit unspecific signal according to the manufacturer’s recommendations (GeneTex, Euromedex, Souffelweyersheim, France). In all cases, immunodetection was performed using the ECL kit according to the manufacturer’s instructions (GE Healthcare, Dutscher, Brumath, France), with an Alliance 9.7-89 EPI apparatus (UVITEC Cambridge, Dutscher) and optical density measured using the imageJ software (NIH). All optical density plots represent normalized values to loading controls, as indicated in the figure legends.

### Immunocytochemistry and pulse labelling

To perform immunocytochemistry, 6.10^4^ HEK cells/well were plated on coverslips for 24 h and then transfected for 48 h with control pcDNA3.1 or plasmids encoding the MT5-MMP variants described above. Endosomes were pulse labelled for 30 min with 0.25 mg/mL AlexaFluor^647^-transferrin (Thermo Fisher Scientific). Cells were then fixed for 15 min with Antigenfix solution (Diapath, MM France, Brignais, France) at room temperature. Immunocytochemistry for all antibodies was performed as follows: after washing in PBS 1X, cells were blocked for 1 h at room temperature using a blocking solution of PBS 1X, 0.1% Triton X-100 and 3% BSA. Cells were then incubated in the blocking solution overnight at 4°C with the primary antibodies APP 6E10 and anti-Aβ40 (Ozyme), anti-GFP (Roche Diagnostics), anti-EEA1 and anti-LAMP-1 (both from Abcam), and anti-FlagM2 (Sigma-Aldrich), followed by appropriate AlexaFluor®-conjugated secondary antibodies for 1 h at room temperature (Thermo Fisher Scientific). For LAMP-1 immunostaining, we used PBS 1X containing 0.1% saponin and 3% BSA. Nuclei were stained with Hoechst (0.5 µg/mL, Thermo Fisher Scientific). Omission of the primary antibody was used as control and no immunostaining was observed. Coverslips were mounted using Prolong Gold Antifading reagent on Superfrost glass slides (Thermo Fisher Scientific). Images were taken and processed using a confocal microscope (LSM 700) and Zen software (Zeiss, Jena, Germany). Co-localization analyses were performed using the Jacop plugin of ImageJ software (NIH) [28].

### Gel zymography

We used gelatin zymography on HEKswe cell supernatants to detect the levels of MMP-2 and MMP-9, also known as gelatinase A and B, respectively. Equal amounts of serum-free supernatants in non-denaturing and non-reducing conditions were subjected to zymogel according to the manufacturer’s recommendations (Thermo Fisher Scientific). Gels were digitized using GeneTools software.

### ELISA assay for Aβ

Levels of Aβ40 in supernatants of HEK or HEKswe cells transfected with different MT5-MMP variants and/or C99 constructs or treated with C3 or DAPT were evaluated by ELISA (#KHB3481, Thermo Fisher Scientific) according to the manufacturer’s recommendations.

### Statistics

All values represent the means ± SEM of the number of independent cultures indicated in the figure legends. ANOVA followed by a Fisher’s LSD *post hoc* test was used to compare more than 2 groups. The Student t-test was used to compare 2 groups. Statistical significance was set up to p<0.05. Analyses were performed with the GraphPad Prism software (San Diego, California USA, www.graphpad.com).

## RESULTS

### MT5-MMP processing of APP generates the sAPP95 fragment independently of other proteolytic activities

To investigate the mode of action of MT5-MMP on APP processing, we first asked how this would compare to that of its close homologue, MT1-MMP, which generates a soluble APP N-terminal fragment of 95 kDa (sAPP95) in concert with endogenous MMP-2 in HEKswe cells [24]. Inactive pro-MMP-2 can be converted to the lower molecular weight active form by MT1- and MT5-MMP upon proteolytic removal of its pro-domain [29,30]. As illustrated by gelatin zymography, MT5-MMP (MT5) catalysed the activation of pro-MMP-2, but much less efficiently than MT1-MMP (Fig. 1A). As expected, the inactive MT5-MMP mutant (MT5Δ) failed to activate pro-MMP-2 (Fig. 1A). In this context, we asked whether MT5-MMP could also require MMP-2 activity to process APP, but first we confirmed that only MT5 and not MT5Δ released sAPP95 (Fig. 1B top panel and C). MT5 did not change the levels of soluble full length APP (sAPPFL) detected with the 22C11 antibody that recognizes the N-terminal domain of APP (Fig. 1B, top panel). On the contrary, MT5 but not MT5Δ significantly decreased the content of sAPPα-like, as shown by the 6E10 antibody, which recognizes the N-terminal sequence of human Aβ and thus, the C-terminal moiety of sAPPα-like fragments (Fig. 1B middle panel and C). We evaluated next the functional interactions of MT5-MMP with MMP-2 and found that a chemical MMP-2 inhibitor did not prevent sAPP95 generation by MT5-MMP (Fig. 1D). A similar result was obtained with TIMP-1, a potent inhibitor of MMP-2 that spares MT5-MMP (Fig. 1E), while TIMP-2, which efficiently inhibits both MMP-2 and MT5-MMP, completely blocked sAPP95 release (Fig. 1F). Together, and contrary to MT1-MMP, these data demonstrate no implication of MMP-2 in the processing of APP by MT5-MMP. To test the influence of canonical secretases in MT5-MMP processing of APP, we used C3 and DAPT, respectively inhibitors of β-secretase BACE1 (Fig. 1G) and γ-secretase (Fig. 1H), as well as a MMP selective pseudophosphinic MMP inhibitor RxP03 [22,31]. Only RxP03 significantly inhibited the release of sAPP95 (Fig. 1I), further confirming that MT5-MMP does not require the assistance of canonical secretases to cleave APP. As the MT5-MMP ectodomain can be physiologically shed from the membrane, we tested if the soluble recombinant catalytic domain of MT5-MMP (cdMT5) inserted in the ectodomain could process sAPPFL into sAPP95. After incubating the cdMT5 for 5 min or 60 min in cell supernatants enriched in sAPPFL, we barely detected any sAPP95 as opposed to the control condition with MT5-MMP-expressing cells. Instead, the cdMT5 completely degraded sAPPFL after 18 h (1080 min) of incubation (Fig. 1J), suggested the need for a cellular environment to finely tune MT5-MMP proteolysis.

### MT5-MMP pro-amyloidogenic features involve proteolytic- and non-proteolytic dependent pathways

After confirmation of our previous work [12] showing that MT5-MMP stimulated the accumulation of Aβ (Fig. 2A and B), we tested whether this could involve canonical β- and γ-secretases, using their respective C3 and DAPT inhibitors. Both prevented the increase in Aβ levels caused by MT5-MMP expression (Fig. 2A and B).

Considering that some MMPs regulate transcriptional activity [32], we tested next whether MT5-MMP could control the expression of genes encoding proteins involved in APP metabolism (Table 2). We studied *MMP24* and *MMP14*, encoding MT5-MMP and MT1-MMP respectively, *APP, BACE1* as well as genes encoding the members of the γ-secretase complex presenilin 1 and 2 (*PSEN1* and *PSEN2)*, nicastrin (*NCSTN*), presenilin enhancer 2 homolog (*PSENEN*) and Aph1a (*APH1A*). We also investigated possible changes in the expression of genes encoding Aβ-degrading enzymes such as *MMP2, MMP9, MME* (neprilysin) and *CTSD* (cathepsin D). We found no significant changes for all genes, except for the expected increase of *MMP24* levels after transfections of the active and inactive variants (Table 2).

**Table 2.** MT5-MMP overexpression does not affect the expression of genes potentially involved in APP/Aβ metabolism. *** p<0.001 *vs* GFP. ANOVA followed by *post hoc* Fisher’s LSD test. n.s.: no significance.

| Genes | Mean (+/- SEM) |  |  | Statistics |
| --- | --- | --- | --- | --- |
| | GFP | MT5 | MT5Δ | |
| *APP* | 1.01 (+/- 0.02) | 0.971 (+/- 0.07) | 0.933 +/- 0.06) | ns |
| *MMP2* | 1.06 (+/- 0.21) | 1.091 (+/- 0.14) | 1.31 (+/- 0.16) | ns |
| *MMP9* | 1.071 (+/- 0.23) | 0.916 (+/- 0.12) | 1.086 (+/- 0.09) | ns |
| *MMP14* | 1.043 (+/- 0.15) | 1.413 (+/- 0.2) | 0.814 (+/- 0.12) | ns |
| *MMP24* | 1.08 (+/- 0.25) | 192.684 (+/- 35.7) | 88.603 (+/- 15.51) | **** |
| *TIMP1* | 1.095 (+/- 0.22) | 0.974 (+/- 0.07) | 1.248 (+/- 0.18) | ns |
| *MME* | 1.061 (+/- 0.19) | 1.162 (+/- 0.15) | 1.188 (+/- 0.25) | ns |
| *CTSD* | 1.006 (+/- 0.07) | 0.989 (+/- 0.06) | 1.082 (+/- 0.17) | ns |
| *BACE1* | 1.084 (+/- 0.23) | 1.196 (+/- 0.13) | 1.247 (+/- 0.15) | ns |
| *NCTSN* | 1.003 (+/- 0.04) | 0.971 (+/- 0.09) | 1.119 (+/- 0.096) | ns |
| *PSENEN* | 1.01 (+/- 0.086) | 0.869 (+/- 0.07) | 1.102 (+/- 0.048) | ns |
| *APH1A* | 1.059 (+/- 0.18) | 0.997 (+/- 0.24) | 1.106 (+/- 0.14) | ns |
| *PSEN1* | 1.044 (+/- 0.16) | 0.852 (+/- 0.11) | 1.017 (+/- 0.049) | ns |
| *PSEN2* | 1.040 (+/- 0.16) | 0.926 (+/- 0.13) | 1.341 (+/- 0.17) | ns |

The above data ruled out a possible transcriptional role of MT5-MMP and left proteolysis as the main putative driver of MT5-MMP pro-amyloidogenic effects. Surprisingly, however both active (MT5) and inactive (MT5Δ) variants significantly increased the extracellular levels of Aβ40 (Fig. 2C), suggesting that the pro-amyloidogenic effects of MT5-MMP do not exclusively depend on its proteolytic activity. This is consistent with our previous data indicating that MT5-MMP modulates APP metabolism by regulating its traffic [20]. In support of this idea, MT5 and MT5Δ significantly increased the content of APP/Aβ immunoreactivity in early endosomes, as shown by combined AlexaFluor^647^-transferrin pulse labelling and immunofluorescence with the 6E10 antibody (Fig. 3A). These results were further confirmed by increased co-localization of an antibody directed against the C-terminal end of Aβ40 and the early endosome marker EEA1 (Fig. 3B) in the absence of changes in the content of endosomes (Supplemental figure 1A). Moreover, we tested the impact of MT5-MMP on the trafficking of 6E10^+^ APP fragments, including Aβ, to the lysosomal degrading pathway. Interestingly, LAMP1^+^ lysosomes from MT5Δ-expressing cells showed a 127% and 96% significant increase in 6E10^+^ immunoreactivity compared to GFP and MT5 groups, respectively (Fig. 3C), while lysosome content increased only by 10% in MT5Δ cells compared to GFP control (Supplemental figure 1B).

We deduce from these data that MT5-MMP could influence APP endo-lysosomal APP traffic to stimulate Aβ production in concert with β- and γ-secretases, and that non-catalytic MT5-MMP domains are likely involved in such a mechanism.

### Differential impact of MT5-MMP domains on APP metabolism

To determine the specific impact of MT5-MMP domains on APP metabolism, we designed plasmid constructs N-terminally tagged with FlagM2 (FlagM2^+^), coding for full length (FL) active (MT5^FL^) and inactive (MT5^Δ^) MT5-MMP, as well as truncated MT5-MMP variants lacking functional domains as follows: catalytic (MT5^CAT-^), hemopexin (MT5^HPX-^), extracellular (MT5^EXT-^), transmembrane and intracytoplasmic domains (MT5^TM/IC-^), and intracytoplasmic domain alone (MT5^IC-^) (Fig. 4A). After transient transfection in HEKswe cells (Fig. 4B, top panel), none of the variants affected canonical intracellular APPFL (Fig. 4B 2^nd^ panel and C) or sAPPFL (Fig. 4B 3^rd^ panel and D). As expected, MT5^FL^ generated sAPP95, while MT5^Δ^, MT5^CAT-^ and MT5 ^EXT–^ did not (Fig. 4B, 3^rd^ panel and E). However and most interestingly, the ability of MT5-MMP to produce sAPP95 was significantly reduced upon deletion of the non-catalytic MT5^HPX-^, MT5^TM/IC-^ and MT5^IC-^ domains (Fig. 4B, 3^rd^ panel and E). We then tested how the expression of MT5-MMP variants with impaired ability to process APP would affect Aβ levels. Among the truncated variants, only MT5^EXT-^ significantly increased the extracellular Aβ40 concentration measured by ELISA (Fig. 4F), indicating that the C-terminal part of MT5-MMP could promote amyloidogenesis on its own.

### The C-terminal domain of MT5-MMP plays a specific role in the control of C99 fate

We next investigated how MT5-MMP could be functionally related to the immediate precursor of Aβ, C99, considering that it is drastically downregulated in 5xFAD mice deficient for MT5-MMP [12], and is increasingly considered to be a relevant neurotoxic factor in AD [8,11,15,14,33]. To test a possible functional interplay between C99 and one or several domains of MT5-MMP, we first mimicked AD-like C99 accumulation by transiently overexpressing a C99 construct in HEK cells. After transfection of the pcDNA3.1 control plasmid, the APP-CTF antibody revealed a ∼13 kDa immunoreactive band matching the predicted molecular weight of C99. A second band of ∼11 kDa (Fig. 5A top panel) was abolished by the α-secretase inhibitor GI254023X (not shown) and was not detected by the 6E10 antibody that recognizes an epitope in the N-terminal portion of Aβ upstream the α-cleavage site (Fig. 5A middle panel). Based on these data, we concluded that the ∼11 kDa band represented C83 resulting from the cleavage of C99 by α-secretase as previously observed [26]. It has to be noted that co-transfection of C99 and the MT5-MMP variants had no effect on cell viability as measured by the MTT test (supplemental figure 2A). In these conditions, we observed a striking reduction of C99 levels in cells expressing MT5^IC-^ and even more in MT5^TM/IC-^ expressing cells (Fig. 5A-C). In contrast, the expression of MT5^HPX-^ and MT5^EXT-^ significantly increased C99 levels when evaluated with the 6E10 antibody (Fig. 5A and C). No difference in C99 levels was observed after transfection of MT5 ^FL^, MT5^Δ^ CAT- and MT5 ^CAT-^. Noticeably, all the MT5-MMP variants except MT5^HPX-^ nearly abolished C83 levels (Fig. 5A and D). ELISA revealed a significant reduction of more than 50% of Aβ40 levels after expression of MT5^TM/IC-^ and MT5^IC-^ (Fig. 5E), implying that the drop in C99 levels caused by these variants was not due to an increased conversion of C99 to Aβ. Of note, no differences in Aβ levels were observed after expression of MT5^HPX-^ and MT5^EXT-^, suggesting that their apparent cumulative effects on C99 levels did not impact Aβ production.

**Figure 5.**
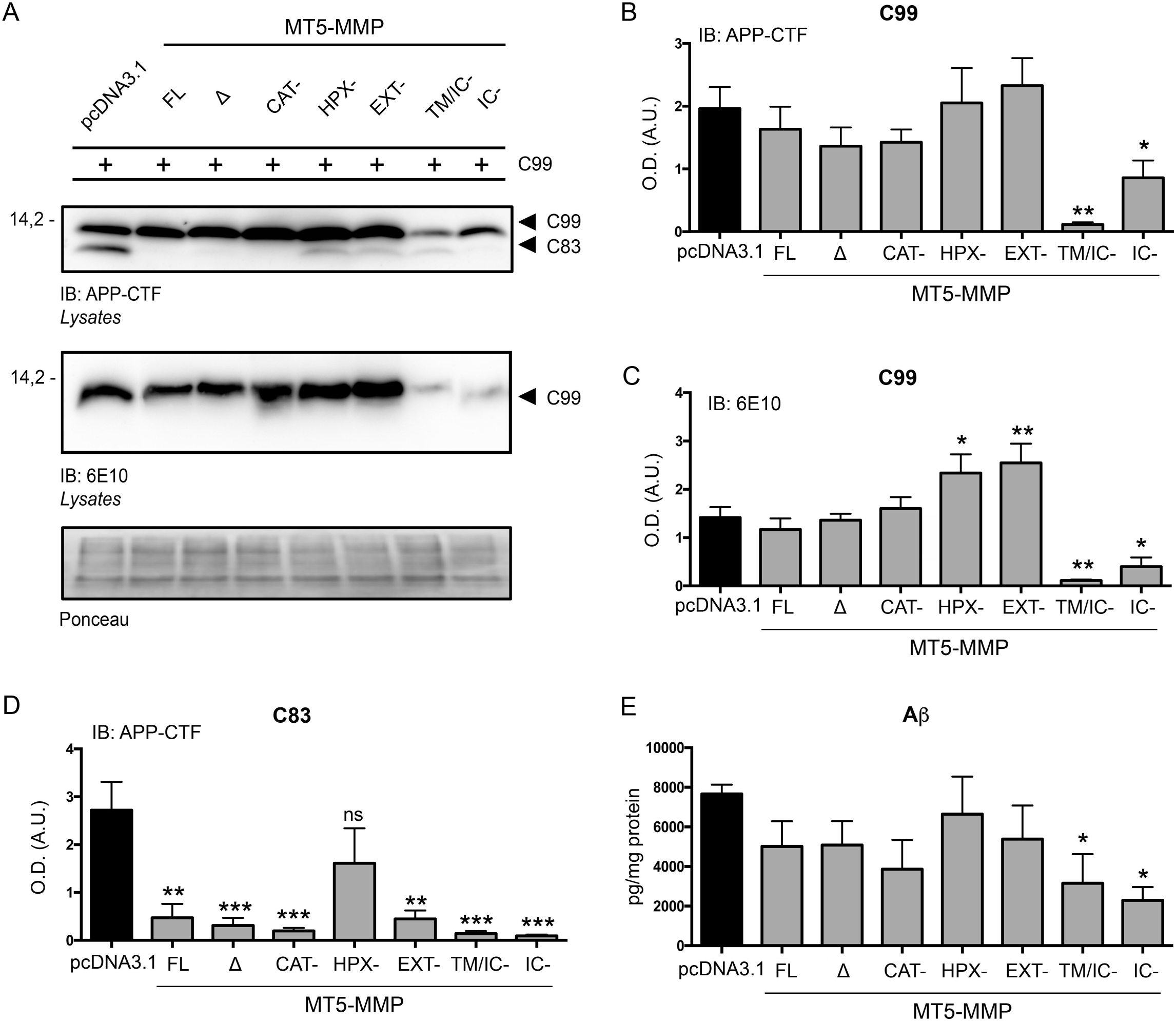
MT5-MMP domains differently affect C99 stability. **A**. Representative WB of C99 and C83 levels in HEK cells 48 h after co-transfection with C99 and either a control pcDNA3.1 plasmid and full length (FL) active or inactive (Δ) MT5-MMP or MT5-MMP without catalytic (CAT-), hemopexin (HPX^-^), extracellular (EXT-), transmembrane/intracytoplasmic (TM/IC^-^) and intracytoplasmic (IC-) domains, using APP-CTF antibody (top panel) and 6E10 antibody (middle panel). The bottom panel represents the loading controls with ponceau. **B, C** and **D**. Histograms represent ponceau-normalized quantifications of C99 detected with both the APP-CTF (B) and 6E10 (C) antibodies, and C83 detected with APP-CTF antibody (D). **E**. Histogram showing ELISA quantification of Aβ40 levels (pg/mg protein) in the supernatants of HEK cells (n=4) 48 h after co-transfection of C99 with either pcDNA3.1, FL, Δ, CAT-, HPX-, EXT-, TM/IC- and IC-. Values are the mean +/-SEM of four independent cultures. * p<0.05, ** p<0.01, *** p<0.001 *vs* pcDNA3.1. ANOVA followed by *post hoc* Fisher’s LSD test. n.s.: no significant.

### The C-terminal domains of MT5-MMP and MT1-MMP have opposite effects on C99 and C83

We wondered next whether the striking drop in C99 levels caused by the deletion of the MT5-MMP C-terminal domains was specific for this proteinase or, instead, whether it shared common features with its close homologue MT1-MMP. Accordingly, we transiently co-expressed C99 and previously described [25] full length active (MT1 ^FL^) and inactive (MT1^Δ^) MT1-MMP, as well as the MT1^IC-^ variant (Fig. 6A). Using the APP-CTF antibody (Fig. 6B) we found that MT1^FL^ and MT1^Δ^ variants did not affect C99 levels compared to pcDNA3.1. On the contrary, the MT1^IC-^ variant caused significant increases of 159% in C99 (Fig. 6B and C) and a 207% in C83 (Fig. 6B and D). These data were in clear contrast with the strong reduction of C99 and C83 levels induced by MT5^IC-^ (Fig. 5D). From the above, we conclude that the intracellular domains of MT5-MMP and MT1-MMP play opposite roles in controlling the fate of C99 and C83, thus revealing their respective specificities in terms of functional interactions with APP.

**Figure 6.**
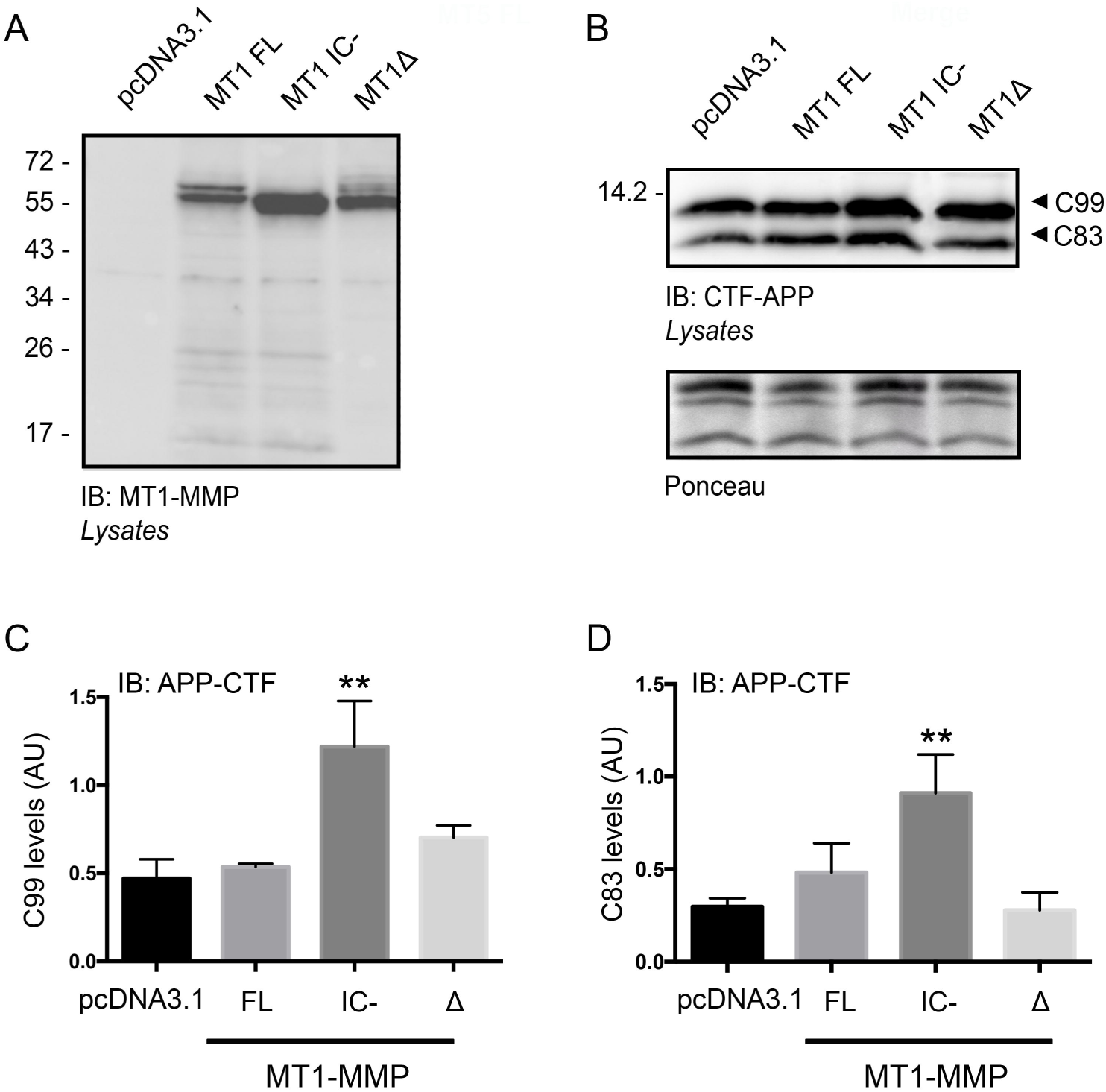
Deleting the intracellular domain of MT1-MMP promotes the accumulation of C99 and C83. **A**. Representative WB of MT1-MMP immunoreactivity in HEK cells revealed by an anti-MT1-MMP antibody 48 h after transfection of pcDNA3.1, MT1-MMP full length (FL), MT1-MMP lacking the intracytoplasmic domain (IC-) and inactive MT1-MMP (MT1Δ). **B**. WB representing C99 and C83 immunoreactivity using an APP-CTF antibody and their respective ponceau-normalized quantifications (**C** and **D**). Values are the mean +/-SEM of four independent cultures. ** p<0.01 *vs* pcDNA3.1. ANOVA followed by *post hoc* Fisher’s LSD test.

### The C-terminal part of MT5-MMP interacts with C99

Considering the major effects of the TM and IC domains of MT5-MMP on C99 levels and our previous results showing interaction of MT5-MMP and APP [12], we looked for specific interactions of the C-terminal part of MT5-MMP that could underlie its effects on C99. Accordingly, we conducted co-immunoprecipitation experiments in HEK cells co-transfected with FlagM2^+^ MT5^FL^ or MT5^EXT-^ variants together with a HA-C99 construct that allowed us to detect transfected C99 from endogenous APP species. HA-C99 (Fig 7A top panel) and MT5-MMP variants (Fig. 7A middle panel) were readily expressed after transfection (inputs). C83 was not detected in this case because the N-terminal HA-tag was lost upon cleavage of C99 by α-secretase. FlagM2^+^ immunoprecipitated complexes were immunoreactive for the HA antibody, demonstrating that the C-terminal domain of MT5-MMP interacts with C99 (Fig. 7B). Unspecific IgG did not immunoprecipitate MT5 constructs (Fig. 7B). Reverse immunoprecipitation with the anti-HA antibody followed by WB with an anti-flagM2 revealed that MT5^EXT-^ and MT5^FL^ were also immunoprecipitated (Fig. 7C). These data demonstrated that TM and/or IC domains of MT5-MMP could mediate C99 interactions themselves.

**Figure 7.**
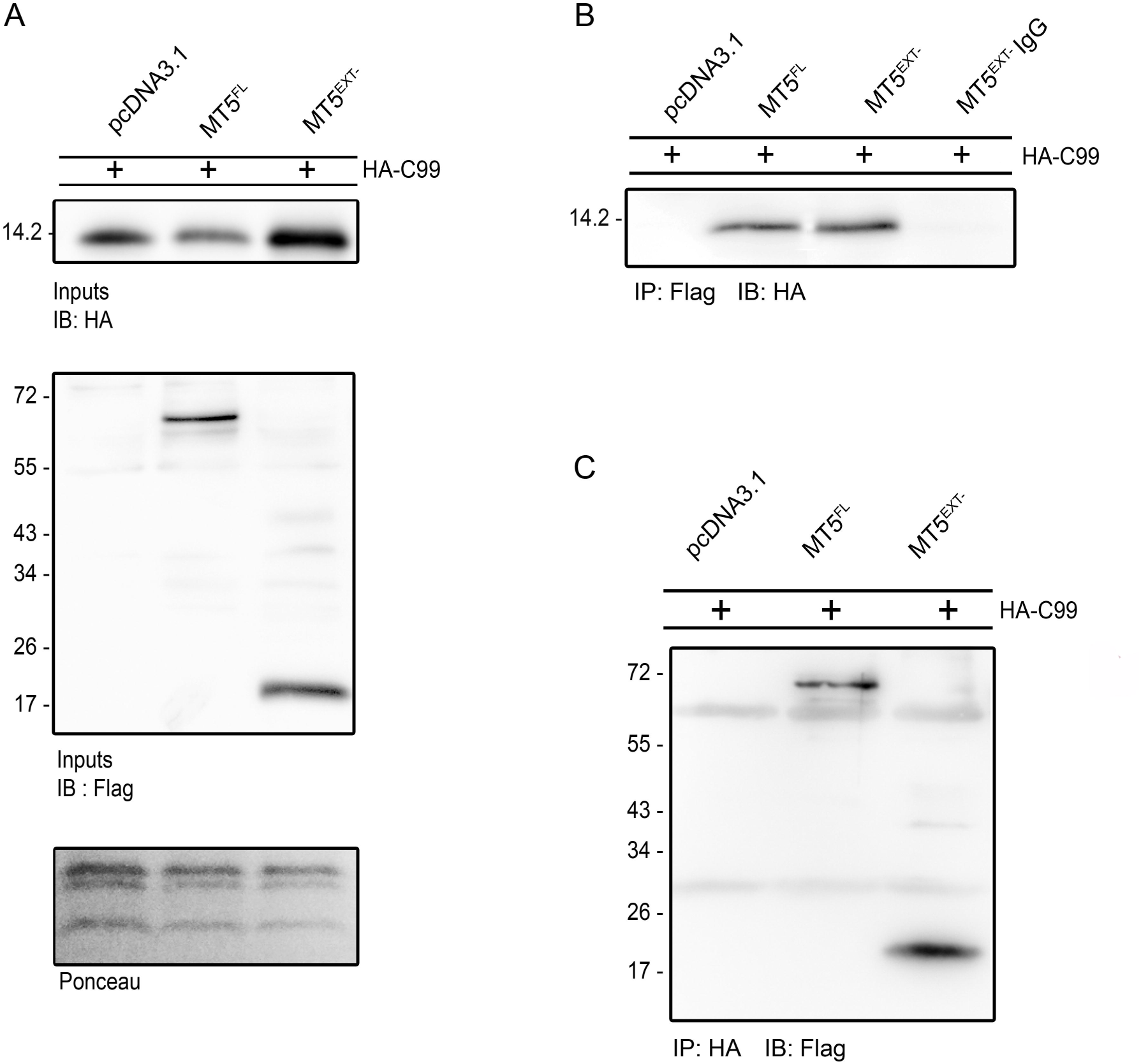
The C-terminal domains of MT5-MMP mediate interactions with C99. **A**. WB showing anti-HA immunoreactivity in HEK cells 48 h after co-transfection with HA-C99 and either pcDNA3.1 control, MT5^FL^ and MT5^EXT-^ constructs (top panel). The middle panel shows representative anti-FlagM2 immunoreactivity for the transfected MT5-MMP variants (input), and the bottom panel, the ponceau loading controls. **B**. WB showing immunoprecipitation (IP) with a FlagM2 antibody or a non-specific IgG as negative control and immunoblot (IB) with an anti-HA antibody. **C**. WB showing reverse IP with an anti-HA antibody and IB with a FlagM2 antibody. Note that both MT5-MMP constructs immunoprecipitated HA-C99, suggesting the implication of the C-terminal domain of MT5-MMP in interactions with C99.

### The C-terminal part of MT5-MMP promotes C99 endosomal sorting

We also wanted to gain insight into the cellular distribution of HA-C99 after co-transfections with different MT5 variants. We first confirmed by WB that MT5^TM/IC-^ reduced the levels of HA-C99 and C83 (Fig. 8A), as previously shown after transfection of non-tagged C99 (Fig. 5A-C). Immunofluorescence experiments after co-transfection with HA-C99 and either MT5^FL^, MT5^EXT-^, MT5^TM/IC-^ or pcDNA3.1, revealed that MT5^EXT-^ was the only variant that significantly increased localization of HA-C99 with the EEA1 antibody, a marker of early endosomes (Fig. 8B-E), while MT5^FL^ and MT5^TM/IC-^ showed values close to pcDNA3.1 control. All the experimental groups exhibited equivalent EEA1^+^ signal, implying that MT5-MMP variants did not alter the content of endosomes (Supplemental figure 2B). Overall, these data further suggest that the C-terminal domains of MT5-MMP modulate the stability and cellular distribution of C99, in particular upon the loss of the extracellular domain of the proteinase.

**Figure 8.**
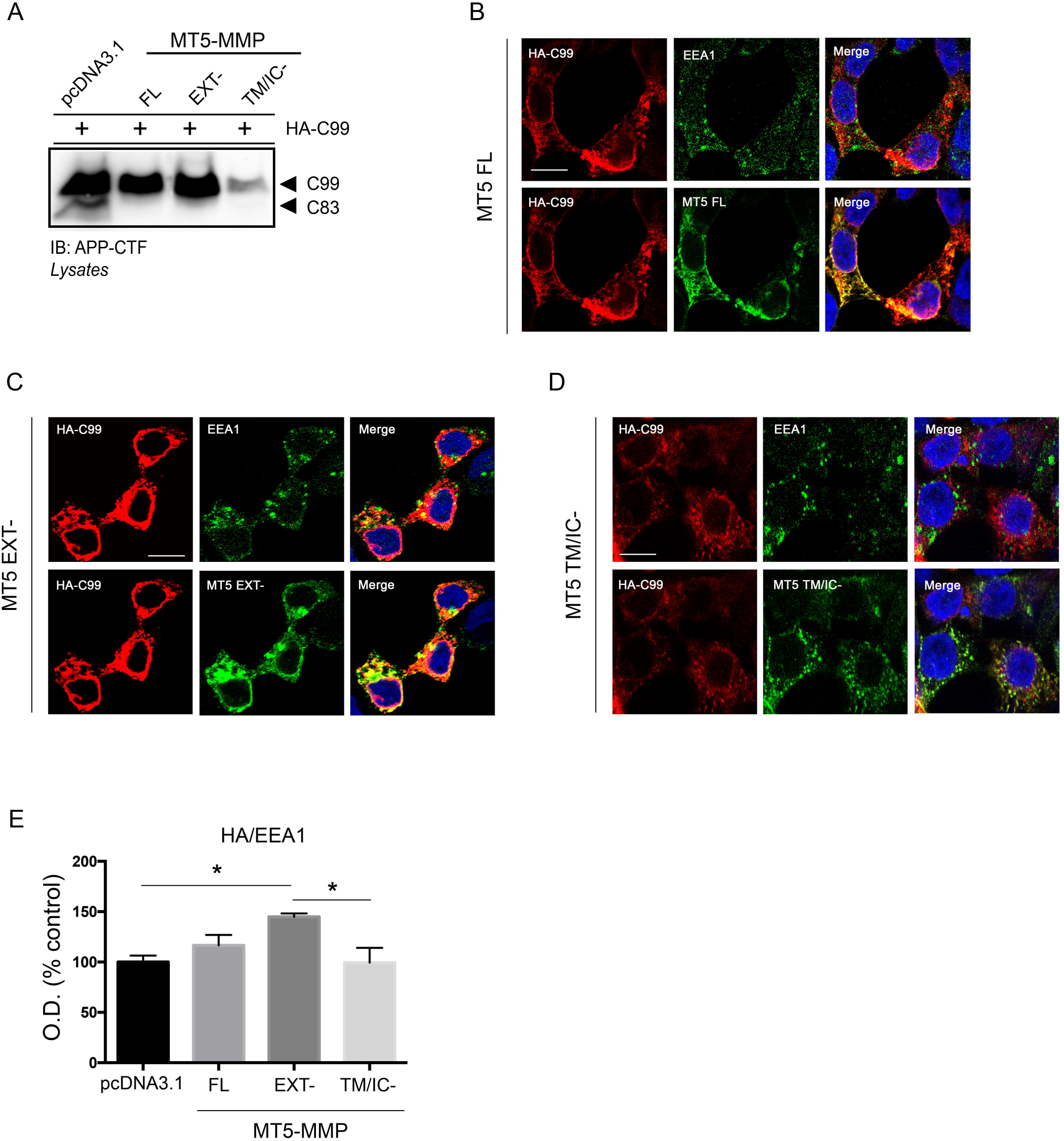
MT5-MMP domains differently affect subcellular C99 localization. **A**. Representative WB of C99 and C83 levels in HEK cells 48 h after co-transfection with HA-C99 and either control pcDNA3.1 plasmid, MT5-MMP full-length (FL) or MT5-MMP lacking the extracellular (EXT-) or transmembrane/intracytoplasmic (TM/IC-) domains, using an APP-CTF antibody. **B, C** and **D**. Confocal micrographs showing combined immunofluorescence with anti-FlagM2 antibody (green), anti-HA antibody (red) and early endosomal marker EEA1 (green) on HEK cells co-transfected for 16 h with HA-C99 and either MT5 FL (**B**), MT5 EXT- (**C**) or MT5 TM/IC-(**D**) constructs. Note that EEA1 antibody was revealed in the same cells with an AlexaFluor^647^ secondary antibody, but for the sake of a sharper contrast with red color, green pseudo-color was used for EEA1 labelling as well. Nuclei stained with Hoechst are in blue. **E**. Histogram showing colocalization between EEA1 and HA-C99 referred as % of pcDNA3.1 control. Only MT5 EXT- caused a significant increase in colocalization between HA-C99^+^ and EEA1^+^ signals. Values represent the mean +/-SEM of four independent cultures. * p<0.05, ANOVA followed by *post hoc* Fisher’s LSD test. Scale bar, 10 µm.

### The control of C99 and C83 fate by MT5-MMP involves different cell scavenging systems

The concomitant reduction in C99 and Aβ levels in cells expressing MT5^TM/IC-^ and MT5^IC-^ (Fig. 5) suggests that C99 is degraded by one or several scavenging systems. In order to further explore this possibility, cells co-expressing C99 with either control pcDNA3.1 or MT5-MMP variants were incubated with DMSO or inhibitors of proteolytic systems that could degrade C99 such as DAPT, bafilomycin A1 (bafilomycin) and MG132, which respectively inhibit the γ-secretase, autophagolysosomal and proteasome activities. Using the APP-CTF antibody on homogenates of cells co-transfected with pcDNA3.1 and C99, we observed that only MG132 induced a significant 66% accumulation of C99 compared to the DMSO control condition, while DAPT and bafilomycin had no effect (Fig. 9A-E). Therefore, in the absence of MT5-MMP, the excess of C99 in HEK cells was primarily eliminated by the proteasome and not by γ-secretase or the autophagolysosomal activities. In contrast with C99, C83 was scavenged not only by the proteasome, but also by γ-secretase. In pcDNA3.1-transfected cells, DAPT and MG132 respectively increased C83 levels by 104% and 83%, while bafilomycin had no effect (Fig. 9A-D and F).

**Figure 9.**
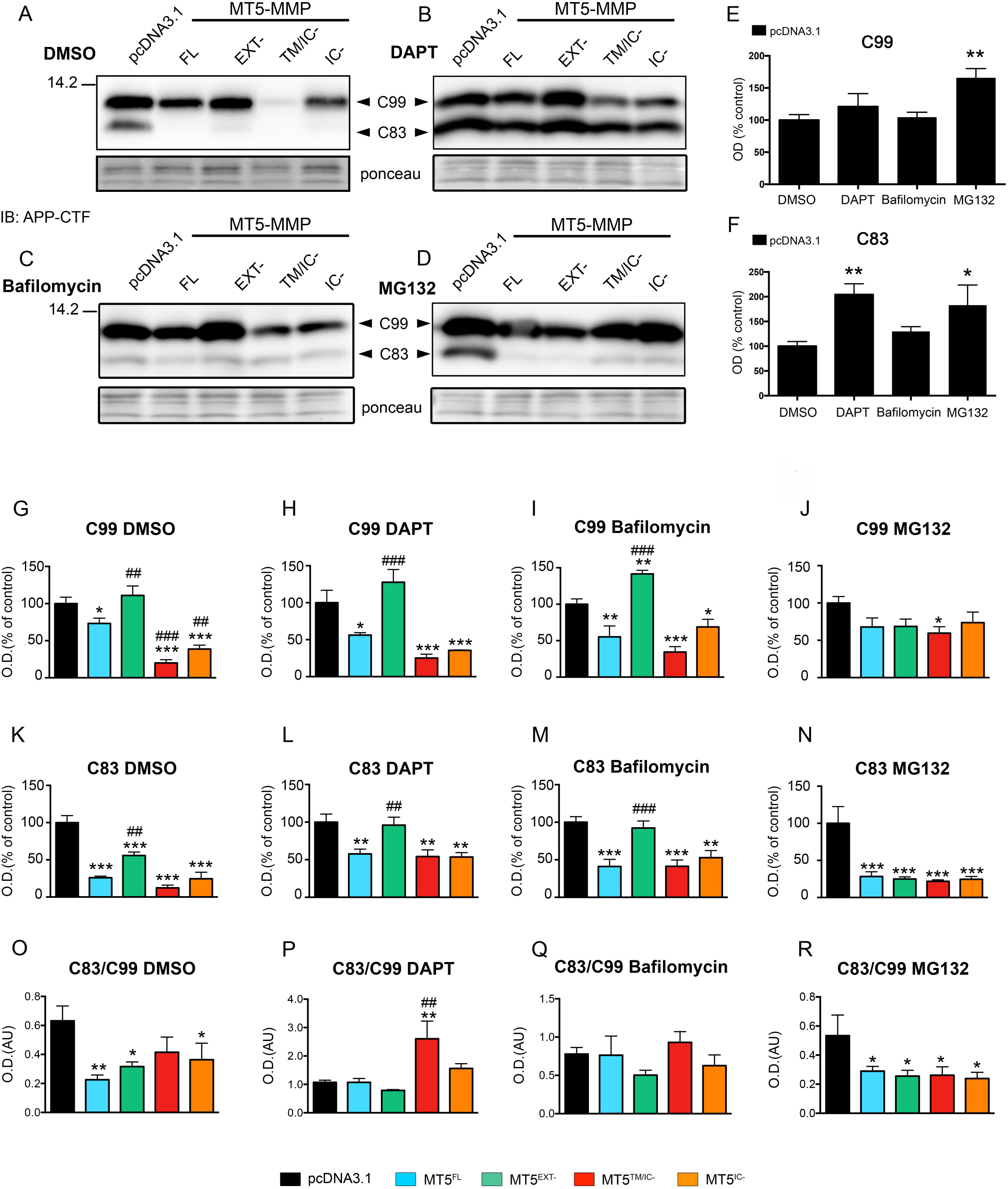
MT5-MMP domains differently affect the fate of C99 and C83, and this is modulated by different scavenging systems. **A, B, C** and **D**. WB showing representative anti-APP-CTF immunoreactivity for C99 and C83 in HEK cells after co-transfection of C99 with either pcDNA3.1, MT5 FL, MT5 EXT-, MT5 TM/IC- or MT5 IC-constructs, and treated or not with DMSO control, DAPT (10 μM for 24 h), Bafilomycin A1 (1 mM for 16 h) and MG132 (5 μM for 24 h). Representative ponceau loading controls are below. **E** and **F**. Histograms respectively representing the quantifications of C99 and C83 levels in pcDNA3.1 transfected cells treated with the different drugs. **G, H, I** and **J**. Histograms representing the quantifications of C99 levels for all MT5-MMP variants relative to pcDNA3.1 control. **K, L, M** and **N**. Histograms representing the quantifications of C83 levels for all MT5-MMP variants relative to pcDNA3.1 control. **O, P, Q** and **R**. Histograms representing the C83/C99 ratio for each experimental group. Values are the mean +/-SEM of four independent cultures. ** p<0.01 *vs* DMSO control (E and F). * p<0.05, ** p<0.01 and *** p<0.001 *vs* pcDNA3.1. ^##^ p<0.01, ^###^ p<0.001 *vs* the MT5 FL group (G-R). ANOVA followed by *post hoc* Fisher’s LSD test.

The expression of MT5-MMP variants deeply affected C99 levels, depending on the pharmacological treatments (Fig. 9A-D and G-J). The most remarkable observation in DMSO treated cells was the drastic decrease of C99 content after expression of MT5^TM/IC-^ (80%) and MT5^IC-^ (64%) compared to pcDNA3.1 (Fig. 9A and G). Likewise, the drop in C99 levels in MT5^TM/IC-^ and MT5^IC-^ cells was also highly significant compared to the MT5^FL^ and MT5^EXT-^ groups. Interestingly, MT5^EXT-^ exhibited 50% higher levels of C99 than MT5^FL^ under DMSO (Fig. 9A and G), and the increase rose to 229% after DAPT treatment (Fig. 9A and H). The difference was even bigger in MT5^EXT-^ expressing cells compared to cells expressing MT5^TM/IC-^ (512%) and MT5^IC-^ (366%), as DAPT failed to rescue the drop of C99 levels in these groups (Fig. 9A and H). Intriguingly, DAPT also accentuated the reduction in MT5^FL^ cells to 44% of pcDNA3.1 C99 values (Fig. 9A and H). Bafilomycin significantly stimulated the accumulation of C99 in MT5^EXT-^ cells with respect to pcDNA3.1 control (41%) and MT5^FL^ (256%). The effect of bafilomycin on the restoration of C99 levels in MT5^TM/IC-^ was barely noticeable when compared with values in DMSO treated cells. Instead, bafilomycin brought C99 levels up to 69% of pcDNA3.1 in MT5^TM/IC-^ cells (Fig. 9C and I). In clear contrast with the aforementioned inhibitors, MG132 significantly restored the levels of C99 in MT5^TM/IC-^ and MT5^IC-^ cells, which reached 59% and 74% of pcDNA3.1 values, respectively (Fig. 9D and J). In the case of the MT5^TM/IC-^ group, the difference was still statistically significant (Fig. 9J). The widespread recovery of C99 levels under MG132 in most groups, together with an apparent lack of effect on MT5^EXT-^ cells, meant that C99 levels for this group were no longer different from those of any other experimental group (Fig. 9J). Overall, the main conclusion is that C99 degradation caused by MT5^TM/IC-^ and/or MT5^IC-^ variants is mainly driven by the proteasome, and to a lesser extent by the autophagolysosomal system, the latter being more efficient in controlling C99 in MT5^EXT-^ cells.

All the MT5-MMP variants caused a drastic reduction of C83 levels under DMSO conditions, ranking from 45% reduction in MT5^EXT-^ cells to 88% in MT5^TM/IC-^ (Fig. 9A-D and K). This is also illustrated in figure 9O, with the generalized decrease of the C83/C99 ratio in this group. DAPT restored C83 levels back to pcDNA3.1 values in MT5^EXT-^ cells. Moreover, DAPT rescued C83 in MT5^FL^, MT5^TM/IC-^ and MT5^IC-^ around 55% of pcDNA3.1 values, clearly above the levels in DMSO control cells (Fig. 9B and L). Figure 9P also illustrates the restoration of C83 by DAPT, as the drug lead to ratio values above 1, except for the MT5^EXT-^ group. The effect of bafilomycin on C83 was collectively less important than that of DAPT (Fig. 9B and M). The main effect of bafilomycin was the restoration of C83 values in MT5^EXT-^ cells compared to pcDNA3.1. In contrast, the levels in the other groups remained low (Fig. 9A, C, K and M). Considering the C83/C99 ratio, bafilomycin was the drug with the mildest effect (Fig. 9Q). MG132 efficiently promoted C83 in cells transfected with pcDNA3.1 (Fig. 9F), but not in those expressing any of the MT5-MMP variants. Their levels were collectively ∼80% lower compared to pcDNA3.1 and roughly unchanged compared to MT5-MMP variants in the DMSO group (Fig. 9A, D, K and N). This was further reflected in the C83/C99 ratio (Fig. 9R). Overall, γ-secretase was the major scavenging system for C83, and the effect was markedly accentuated in cells expressing MT5^EXT-^.

## DISCUSSION

This study provides new evidence that the contribution of MT5-MMP in controlling APP metabolism and accumulation of Aβ and C99 occurs through both proteolysis-dependent and - independent mechanisms. MT5-MMP processing of APP in HEKswe generates a major soluble truncated APP form of 95 kDa (sAPP95), a process that is controlled by the C-terminal domains of the MMP, but does not involve endogenous MMP-2 or canonical secretases. The latter are instead necessary to MT5-MMP pro-amyloidogenic effects. Interestingly, the inactive form of MT5-MMP can also increase Aβ levels, and non-catalytic domains of the MMP interact with C99 and promote its endosomal sorting. The deletion of the C-terminal domains from MT5-MMP and MT1-MMP, revealed the functional specificities for these close homologues, illustrated by the opposite effects they caused on the accumulation of C99, C83 and Aβ. C99 degradation in cells expressing MT5-MMP variants involved mainly the proteasome in the absence of the C-terminal part of the proteinase, whereas the presence of the C-terminal domains diverted the scavenging process to the autophagolysosomal system. C83 was largely scavenged by γ-secretase and to a lesser extent by the autophagosome-lysosome. Overall, these data reinforce MT5-MMP as a pivotal enzyme in APP processing and unveil unexpected and hitherto unseen roles of its C-terminal non-catalytic domains in the control of proteolytic and non-proteolytic events that influence APP processing and the fate of its major toxic C99 and Aβ metabolites.

### MT5-MMP generates a major form of soluble APP in a proteolysis- and cell-dependent manner

We previously reported that MT5-MMP and MT1-MMP stimulate the production of sAPP95 *in vitro* and/or in the brains of 5xFAD mouse model of AD [12,20,24]. Previous studies identified the η-cleavage site for this fragment between N^579^ and M^580^ of APP_770_ (…VLAN^579^-M^580^ISEPR), which is shared by MT1-, MT3- and MT5-MMP [34,35,21]. MMP-2 cooperates with MT1-MMP to release sAPP95 [24], which is consistent with the pro-convertase role of MT1-MMP on MMP-2 [29,36,30]. Although MT5-MMP also activates MMP-2 [29,36,30], our genetic and pharmacological manipulations demonstrated that MMP-2 activity does not cooperate with MT5-MMP in APP processing, possibly due to a relatively ineffective activation of MMP-2 by MT5-MMP, compared to MT1-MMP. Active MT5-MMP did not affect the levels of sAPPFL, but reduced those of sAPPα-like, suggesting a possible competition with α-secretase for the substrate. On the other hand, we cannot exclude a regulatory effect of MT5-MMP on α-secretase activity, such as that reported for metalloproteinase meprin β, which can shed and hence downregulate ADAM10 activity [37].

The recombinant catalytic domain of MT5-MMP failed to generate sAPP95 in a cell-free medium enriched in sAPPFL, probably indicating the need for a cellular microenvironment that provides molecular conformation for fine-tune the specific proteolytic interactions between MT5-MMP and full-length APP. This is consistent with the growing experimental evidence that MMPs exert controlled biological activities in the pericellular and intracellular environment [38].

### The generation of sAPP95 by MT5-MMP does not involve other secretases

The finding that BACE1 inhibition did not affect sAPP95 levels indicates that β-secretase is not involved in the processing of APP by MT5-MMP, in pace with the observation that MT5-MMP does not generate sAPP95 from larger sAPP fragments. In addition, stable sAPP95 levels under BACE1 inhibition suggest that there is no increase in the bioavailability of APP for MT5-MMP. On the other hand, the selective inhibition of MMPs and not ADAMs by RXP03 [22], as well as the suppression of sAPP95 generation by RXP03, also demonstrates that ADAMs (*i*.*e*., α-secretase) do not contribute to this cleavage.

Although we cannot exclude that other recently discovered APP-processing enzymes (*e*.*g*., meprin β, δ-secretase [1,39,2] might cooperate with MT5-MMP, this is rather unlikely because HEK cells do not constitutively express meprin β, and even when overexpressed, this metalloproteinase does not release sAPP95 [40,41]. Moreover, TIMPs do not inhibit meprins [42], while TIMP-2 efficiently blocked the generation of sAPP95 by MT5-MMP. There is also no evidence that TIMPs inhibit the cysteine proteinase δ-secretase.

### Active and inactive MT5-MMP promote Aβ accumulation

One important finding of this work is that inactive MT5-MMP can also promote Aβ accumulation in HEKswe cells. Aβ accumulation in early endosomes of cells expressing MT5Δ/GFP indicates that changes in the trafficking of APP or its metabolites may underlie this event, which confirm previous observations [20]. However, unlike active MT5/GFP, MT5Δ/GFP also promoted the accumulation of APP-6E10^+^ species in lysosomes, indicating an easier sorting of APP species and/or a dysfunction of the lysosomal degradative activities leading eventually to Aβ accumulation. Previous work reported the ability of the metalloproteinase ADAM30 to promote APP sorting in lysosomes by non-catalytic mechanisms. However, unlike MT5-MMP, the action of ADAM30 led to the degradation of APP by cathepsin D, thus preventing the formation of toxic APP catabolites [39]. Moonlighting actions of proteinases may logically require specific interactions of their non-catalytic domains with the target protein. In this line, we found that the removal of the non-catalytic transmembrane domain and the intracytoplasmic tail of MT5-MMP reduced its efficiency in releasing sAPP95, possibly indicating a destabilization of MT5-MMP/APPswe interactions. On the other hand, cells expressing the MT5^EXT-^ variant lacking a catalytic site, unsurprisingly abolished the formation of sAPP95. However, these cells concomitantly accumulated Aβ, suggesting a functional dissociation between MT5-MMP-mediated shedding of the APP ectodomain and intracellular events leading to Aβ generation. This could be physiologically relevant, as MT5^EXT-^ is generated in the trans-Golgi network (TGN) after MT5-MMP shedding by furin [43]. In this scenario, the C-terminal tail of MT5-MMP could be exported from the TGN to the endosomal system with APP or its fragments (*i*.*e*., C99) to facilitate cleavage by β- or γ-secretase and at the same time preclude C99 degradation by the proteasome.

### Functional interactions between the C-terminal domain of MT5-MMP and C99

The finding that C99 overexpression in mice causes AD-like symptoms and pathology [9,10], prompted us to overexpress C99 in HEK cells to determine a possible causal effect between the suppression of the C-terminal part of MT5-MMP and the striking reduction of C99 and Aβ levels. Such reductions are reminiscent of those we described in 5xFAD mice deficient for MT5-MMP [12,20] and highlight the transmembrane/intracellular domains of MT5-MMP key players in the pathogenic actions of the proteinase. This idea is supported by our data demonstrating that the C-terminal part of MT5-MMP interacts with C99, which may in addition assure the interactions of the canonical proteins previously described [12]. Deciphering the functional interactions between MT5-MMP and C99 is consistent with growing evidence implicating both proteins in the pathogenesis of AD [21,12,9,20,15,14].

### The C-terminal domains of MT5-MMP and MT1-MMP hold functional specificity towards C99 and C83

The accumulation of C99 and C83 after deletion of the IC domain of MT1-MMP was the inverse of the effect caused by the deletion of the IC domain of MT5-MMP, strongly suggesting specific effects of the two homologue proteinases. Although they share 55.6% sequence identity, in the short IC domain of 20 amino acids the homology falls to only 20% [2], which might explain their specificities of action. In this context, only the active MT1-MMP, not the inactive one, is pro-amyloidogenic, and may even behave in some cases as a β-secretase-like [24]. In addition, MT5-MMP has a short membrane lifetime compared to MT1-MMP [44], which likely influences the nature of the proteins with which it interacts and thus their trafficking. In any case, the interactions of the IC domains of MT-MMPs with APP add to the paucity of literature on the non-proteolytic functions of these proteinases. For instance, deletion of MT1-MMP IC has been shown to inhibit chemotactic migration of macrophages independently of proteolysis [25], while deletion of the last 3 residues at the carboxyl terminus of MT5-MMP prevents its recycling to the membrane and therefore the ability to activate pro-MMP-2 [44]. The removal of the C-terminus end of MT5-MMP prevents interactions with the PDZ domain containing protein Mint3. It is noteworthy that Mint3 also interacts with APP to promote its export from the Golgi complex into the endo-lysosomal system [45]. Together, these data raise the possibility of an APP/Mint3/MT5-MMP tri-molecular complex that could promote APP/C99 traffic towards endosomes. In favour of this idea, MT5^EXT-^ shows pro-amyloidogenic effects in HEKswe, but it does not when co-expressed with C99 in HEK cells. Such differences could depend on the subcellular distribution of canonical APP and C99, as suggested by a recent study showing that overexpressed C99 is mainly found in the TGN [46]. If the C-terminal part of MT5-MMP contributed to export C99 from the TGN to the endosomal system, its removal could drive C99 into other trafficking/degradation pathways, as suggested below.

### The fate of C99 and C83

Our study shows that the proteasome is a prominent scavenging system for C99 under control conditions, in agreement with previous data also obtained in a HEK model of C99 overexpression [47], and in primary neurons from transgenic mice overexpressing C99 [48].

The impact of MT5-MMP variants was determined by the presence or absence of the C-terminal part of MT5-MMP, as MT5^TM/IC-^ and MT5^IC-^ promoted C99 degradation, whereas MT5^EXT-^ provided stability. Not affected by γ-secretase inhibition, on the contrary, inhibition of the autophagolysosomal pathway promoted C99 accumulation, reinforcing the hypothesis that the C-terminal of MT5-MMP contributes to the endo-lysosomal sorting of C99. Consistent with this, removal of TM/IC or IC domains could interfere with the export of C99, thereby activating the degradation of the endoplasmic reticulum associated protein (ERAD), which involves the activation of the proteasome. In support of this idea, we show that the proteasome inhibitor MG132 is the only drug that significantly restores C99 levels in MT5^TM/IC-^ and MT5^IC-^ cells. Furthermore, it has been shown in a glioma cell line that C99 overexpression triggers its polyubiquitination and subsequent degradation by the proteasome [46].

C83 generated from C99 by α-secretase is readily degraded by γ-secretase in cells expressing MT5-MMP variants, as revealed by DAPT treatment. However, DAPT only causes full recovery of C83 levels in MT5^EXT-^ expressing cells, which argues for more efficient targeting of C83 by the truncated C-terminal domains of MT5-MMP to subcellular compartments with γ-secretase activity. On the other hand, it has been recently shown that α-secretase can process APP in the TGN [49]. Taken together, these data support the possibility that some C99 in the TGN is converted to C83 by α-secretase and that the C-terminal part of MT5-MMP targets both the unconverted C99 and C83 from the TGN to the endo-lysosomal system for further processing. This is coherent with the observation that C99 and C83 accumulate upon inhibition of the endo-lysosomal pathway by bafilomycin. Further support stems from MG132 treatment that prevented the distinct accumulation of C99 observed in cells expressing MT5^EXT-^ compared to other variants under DAPT or bafilomycin. Therefore, the presence of the truncated C-terminal part of MT5-MMP likely prevents the ERAD/proteasome-mediated degradation of C99. The lack of effect of MG132 on C83 suggests that γ-secretase could readily process it in the TGN after α-secretase cleavage of C99. This is plausible considering the existence of α- and γ-secretases molecular complexes that work in concert to process APP [50]. In addition, part of C83 generated at the plasma membrane from the C99 pool that followed the secretory pathway, could undergo endocytosis through the endo-lysosomal pathway. These non-exclusive hypotheses are both consistent with C83 being a preferential substrate of γ-secretase [51,46,52], and places α-secretase as an efficient scavenger of C99 working in concert with γ-secretase.

In summary, the present work provides evidence that different structural domains of MT5-MMP, including non-catalytic, specifically control APP metabolism and the fate of their main C-terminal fragments *i*.*e*., C99, C83 and Aβ. Our findings highlight for the first time non-proteolytic actions of MT5-MMP as a key pathogenic complement to proteolysis, and point out the C-terminal domains of MT5-MMP as possible targets in the control of cellular detoxification in AD.

## ACKNOWLEDGEMENTS

This work was supported by funding from the CNRS and Aix-Marseille Université and by public grants overseen by the French National Research Agency (ANR), MAD5 to SR and FC. The work was also supported by the DHUNE center of excellence and a CoEN grant to SR and EN, by the “Fondation Plan Alzheimer” to SR and EN, by France Alzheimer to SR, by Vaincre l’Alzheimer grants to SR and to EN, by a grant to SR and MK from “Fonds Européens de Développement Régional” FEDER in PACA, by a grant from “La fondation NRJ-Institut de France” to EN, and by grants to FC from “Fondation pour la Recherche Médicale” and by the “Conseil Général des Alpes Maritimes” through the LABEX (excellence laboratory, program investment for the future) DISTALZ (Development of Innovative Strategies for a Transdisciplinary Approach to ALZheimer’s disease). KB was granted a research associate fellowship (Management of Talents) by Excellence Initiative of Aix-Marseille University - A*MIDEX, a French “Investissements d’Avenir” programme” and the “Fondation Plan Alzheimer”. LGG was granted by the ANR and by the “Fondation pour la Recherche Médicale” FRM FDT201904008423. J-MP was recipient of a doctoral fellowship from Vaincre l’Alzheimer. Dominika Pilat is recipient of a funding from Excellence Initiative of Aix-Marseille Université - A*MIDEX, a French “Investissements d’Avenir” program through the Integrative and Clinical PhD program”. RXP03 was kindly provided by Dr. Vincent Dive. MT1-MMP constructs (MT1^FL^, MT1^Δ^ and MT1^IC-^) were kindly provided by Pr. Motoharu Seiki and Pr. Takeharu Sakamoto.

The authors declare no financial conflict of interest that might be construed to influence the results or interpretation of the manuscript.

## Non-standard abbreviations

3xTg: mouse model of Alzheimer’s disease expressing human *APP, PSEN1* and *MAPT* genes with familial mutations
5xFAD: mouse model of Alzheimer’s disease bearing 5 familial mutations on human *APP* and *PSEN1*
Aβ: amyloid beta peptide
AD: Alzheimer’s disease
ADAM: a disintegrin and metalloproteinase
Aη-α: APP fragment generated by η- and α-secretase
APP: amyloid beta precursor protein
BACE: β-secretase, beta-site APP cleaving enzyme 1
C3: BACE-1 inhibitor IV
C99: the last C-terminal 99 aminoacids of APP
CAT: catalytic
CTF: C-terminal fragment
CTSD: cathepsin D
DAPT: (2S)-N-[(3,5-Difluorophenyl)acetyl]-L-alanyl-2-phenyl]glycine 1,1-dimethylethyl ester, γ-secretase inhibitor
EEA1: early endosome marker
FL: full length
GFP: green fluorescent protein
GI254023X: specific ADAM10 inhibitor
HEK: human embryonic kidney cells
HEKswe: HEK carrying the familial Swedish *APP* mutation
HPX: hemopexin
HRP: horseradish peroxidase
IB: immunoblot
IC: intracytoplasmic
IP: immunoprecipitation
iPS: induced pluripotent stem cell
LAMP1: lysosome marker
LTP: long-term potentiation
MG132: D-Leucinamide, N-[(phenylmethoxy)carbonyl]-L-leucyl-N-[(1S)-1-formyl-3-methylbutyl]-, proteasome inhibitor
MME: membrane metallo-endopeptidase, neprilysin
MMP: matrix metalloproteinase
MT5/1-MMP: membrane-type 5/1-matrix metalloproteinase
NCSTN: nicastrin
PSEN1/2: presenilin 1/2
PSENEN: presenilin enhancer 2 homolog
qPCR: quantitative polymerase chain reaction
RXPO3: pseudophosphinic specific inhibitor of MMPs
sAPP95: soluble fragment of APP of 95 kDa
sAPPα: soluble fragment of APP generated by α-secretase
TIMP-1/2: tissue inhibitor of matrix metalloproteinase-1/2
TM: transmembrane

## FIGURE LEGENDS

**Supplemental figure 1**

**A**. Confocal micrographs showing combined GFP fluorescence labelling (green) and AlexaFluor^647^-transferrin (red), used in pulse labelling experiments for 30 min to label early endosomes, 48 h after transfection of HEKswe cells with GFP control, active (MT5) or inactive (MT5Δ) MT5-MMP/GFP. Nuclei are stained with Hoechst (blue). The histogram shows that there is no difference in the % of the area occupied by transferrin labelling after MT5 or MT5Δ expression compared to GFP control. **B**. Histogram showing the % of area occupied by the lysosomes using the anti-LAMP-1 antibody on HEKswe cell after transfection with constructs coding for GFP, MT5 or MT5Δ. * p<0.05 *vs* GFP. ANOVA followed by *post hoc* Fisher’s LSD test.

**Supplemental figure 2**

**A**. Cell viability assessed by MTT assay in HEK cells (n=3), 48 h after co-transfection of C99 with either a control pcDNA3.1 plasmid or full-length active (FL) or inactive (Δ) MT5-MMP or MT5-MMP without catalytic (CAT-), hemopexin (HPX-), extracellular (EXT-), transmembrane/intracytoplasmic (TM/IC-) and intracytoplasmic (IC-) domains. **B**. Histogram showing the % of area occupied by endosomes using the anti-EEA1 antibody on HEK cells after co-transfection of the C99 plasmid with either pcDNA3.1 control, or full-length MT5-MMP (FL) or MT5-MMP without the extracellular (EXT-) or the transmembrane/intracytoplasmic (TM/IC-) domains.

