## Supplementary figures and images for "MT5-MMP controls APP and β-CTF/C99 metabolism through proteolytic-dependent and -independent mechanisms relevant for Alzheimer’s disease"

### Supplemental figure 1

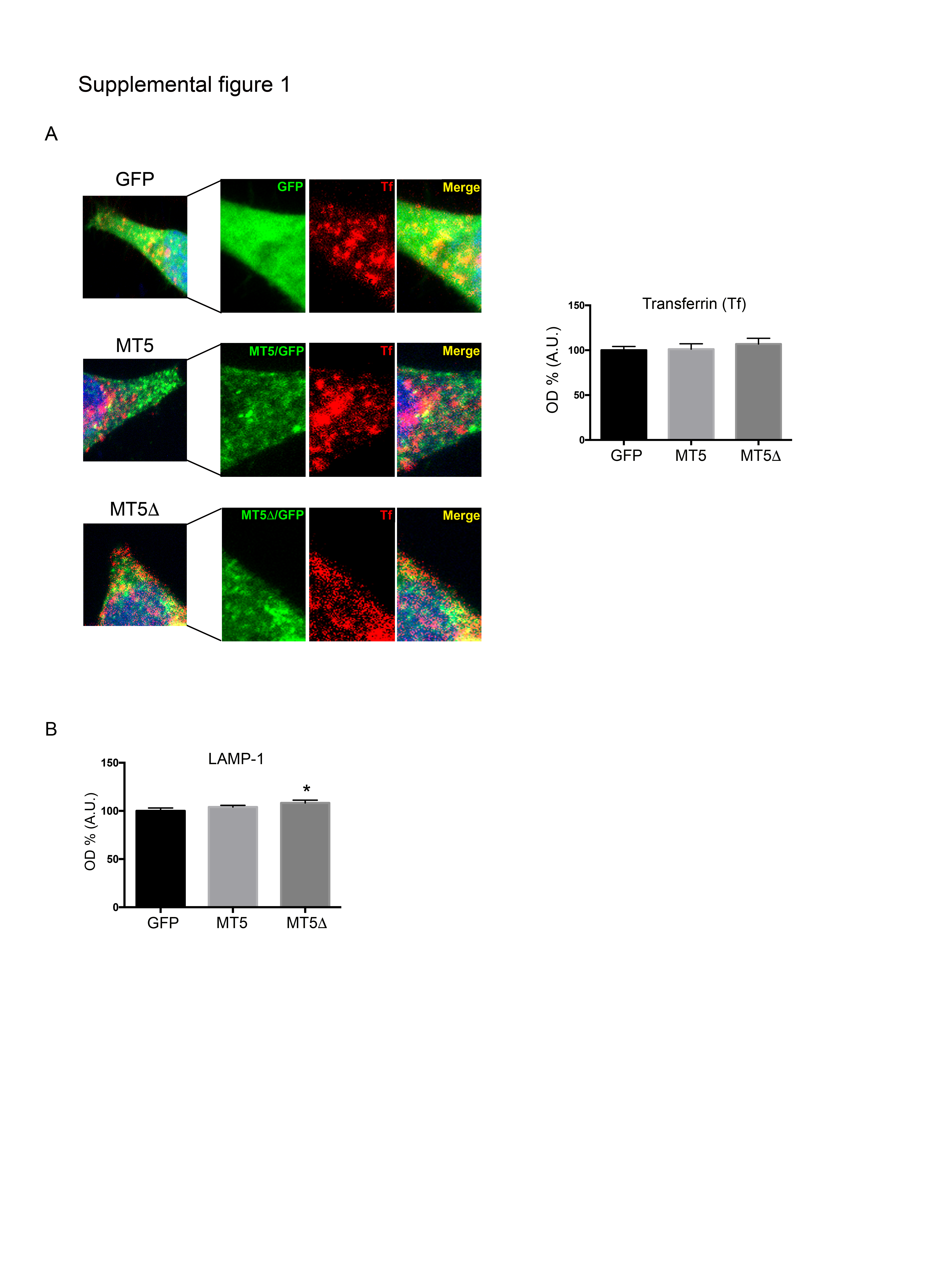

### Supplemental figure 2

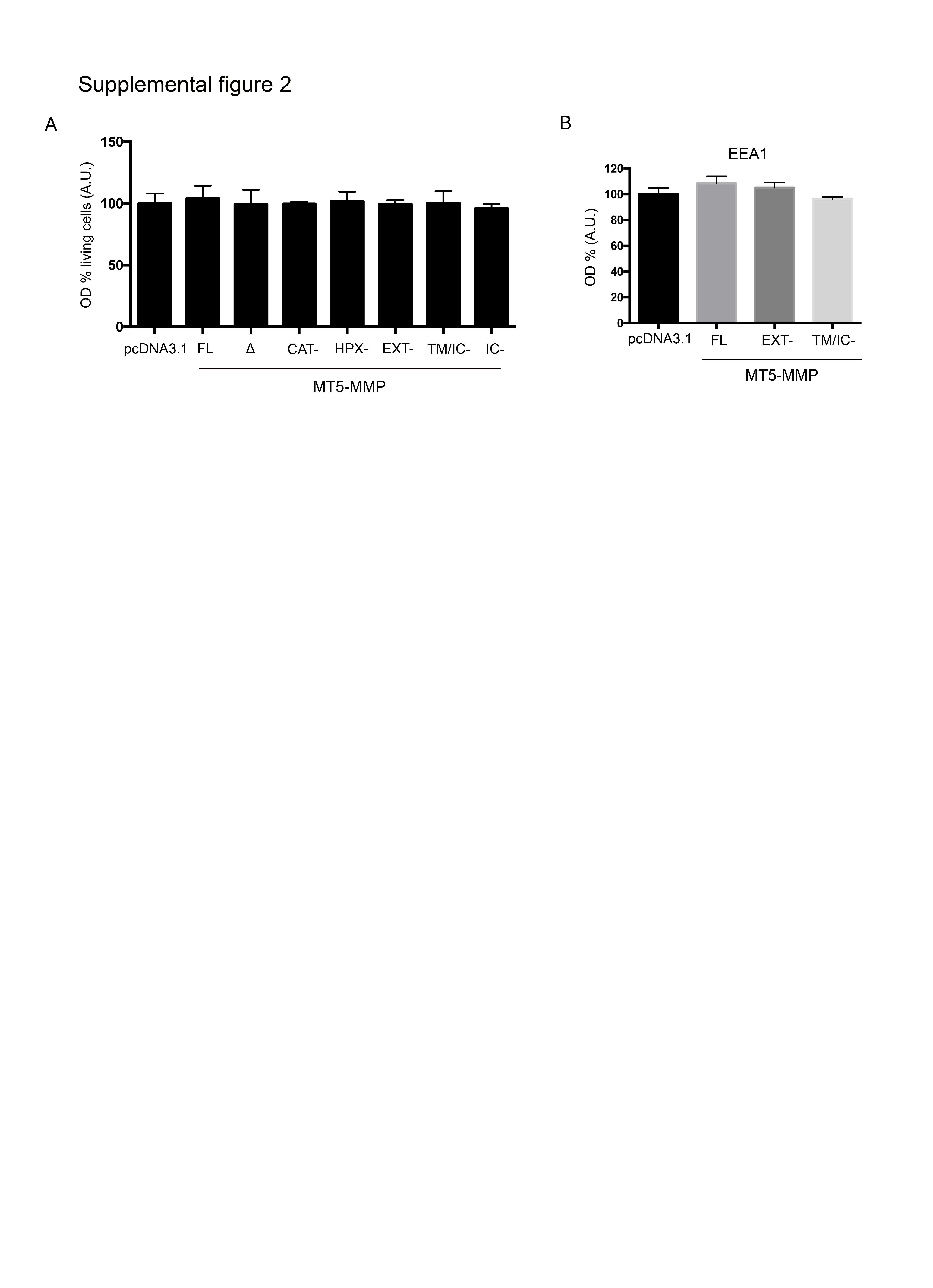
